# Systematic real-time profiling of *Salmonella* type III effector translocation provides quantitative resolution of the T3SS-1/T3SS-2 secretion dichotomy

**DOI:** 10.64898/2026.03.06.710167

**Authors:** Veronique Jonckheere, Laure Simoens, Petra Van Damme

## Abstract

Type III secretion systems (T3SSs) enable *Salmonella* to modulate host cell biology by delivering a diverse repertoire of effector proteins. Yet, the temporal dynamics of effector translocation during prolonged infection remain poorly defined. Here, we establish a systematic real-time framework for quantifying effector secretion and translocation by combining endogenous HiBiT-tagging with split NanoLuc-based detection. Using this approach, we monitored the translocation kinetics of all 39 currently annotated type III effectors (T3Es) in *Salmonella enterica* serovar Typhimurium SL1344 over a 24-hour infection of epithelial cells. This comprehensive analysis revealed temporally distinct translocation patterns, widespread mid- and late-stage effector delivery, and a quantitatively resolved functional overlap between T3SS-1 and T3SS-2-mediated secretion than previously appreciated. These findings provide a more refined view of effector deployment throughout infection and illuminate previously overlooked aspects of effector dynamics. Altogether, this work offers a detailed temporal map of effector dynamics during epithelial infection and establishes a scalable strategy for dissecting secretion system function across host-pathogen systems.

## INTRODUCTION

*Salmonella enterica*, a major cause of gastroenteritis and systemic infections, employs a sophisticated infection strategy to ensure its survival and proliferation within host cells. Central to this process is the injection of approximately 40 type III effector proteins into host cells, which collectively manipulate key host cellular pathways to alter infection outcomes^1, 2^. These effectors modulate diverse processes, including signal transduction, membrane trafficking, cytoskeletal remodeling, and immune responses^1, 2^. Type III effectors (T3Es) are delivered via the type III secretion system (T3SS), a molecular injectisome that mediates ATP-energized secretion^3^ and operates according to a tightly coordinated hierarchy of substrate delivery^4, 5^. A deeper understanding of the complex temporal and quantitative dynamics of effector deployment is therefore essential for elucidating host–pathogen interactions and the mechanisms underlying bacterial pathogenesis.

Several innovative methodologies have advanced our ability to study effector secretion and translocation^6^. Fluorescence-based systems, such as fluorescence resonance energy transfer (FRET), have enabled real-time visualization of effector–host interactions^7^, while enzymatic reporter assays (e.g., β-lactamase–nitrocefin turnover^8^ or calmodulin-activated adenylate cyclase (CyaA)-mediated cAMP accumulation^9^) provide indirect measures of effector injection. Direct fluorescent labeling of effectors, such as tetracysteine-tagged proteins^10^, has enabled live-cell imaging of secretion events, albeit with requirements for complex microscopy setups and limitations in throughput or multiplexing. Collectively, these methods have illuminated important aspects of effector biology, but they remain constrained by sensitivity, substrate availability, perturbation of host physiology, and limited capacity for long-term, quantitative measurements, underscoring the need for routine, scalable, and high-throughput approaches.

The development of split NanoLuc luciferase-based assays by the Wagner laboratory has substantially expanded the ability to quantify effector translocation in real time^11, 12^. By leveraging the expression of HiBiT-tagged effectors in bacteria and complementary LgBiT protein in host cells, interactions between HiBiT and LgBiT in the presence of the NanoLuc substrate allow for the real-time monitoring of effector translocation. This approach enables sensitive, non-invasive, and continuous readout of delivery, thereby facilitating quantitative assessments of effector dynamics throughout bacterial infection.

Studies by Finn *et al*. have shown that interactions between *Salmonella* and epithelial cells persist well beyond initial invasion and depend in part on sustained type III secretion system 1 (T3SS-1) activity^13^. These findings highlight the importance of examining effector translocation over extended infection periods, particularly during the second wave of T3SS-1 activity and the transition to intracellular replication. Recent advances in long-duration NanoLuc substrates, such as vivazine, further facilitate continuous live-cell luminescence detection over many hours, enabling interrogation of otherwise inaccessible late-secretion events. Building on this foundation, we employ a systematic real-time NanoLuc-based framework to comprehensively quantify translocation dynamics for all 39 currently annotated T3Es encoded by *Salmonella enterica* serovar Typhimurium (*S*. Typhimurium) SL1344. By monitoring effector kinetics over 24 hours of epithelial infection and integrating secretion analyses with T3SS-1–deficient (Δ*invA*) and type III secretion system 2 (T3SS-2)– deficient (Δ*ssaV*) strains, we delineate temporal classes of effector delivery and evaluate the relative contributions of both secretion systems. These analyses reveal a broader than previously appreciated overlap in *Salmonella* pathogenicity island 1 (SPI-1)/*Salmonella* pathogenicity island 2 (SPI-2) secretion specificity^14^ and provide a refined view of how *Salmonella* deploys its effector repertoire during distinct infection stages. Our work establishes a scalable strategy for dissecting effector biology and lays a foundation for exploring secretion dynamics across diverse host– pathogen systems^15^.

## RESULTS

### Endogenous HiBiT-tagging enables sensitive detection of the Salmonella effector repertoire under native expression conditions

To systematically quantify type III secretion and translocation dynamics during infection, we generated a comprehensive panel of *S*. Typhimurium SL1344 strains carrying endogenous C-terminal HiBiT fusions to all 39 currently annotated T3Es^2^, including the recently identified SPI-2 effector SseM^16^. This set also includes the T3SS-1 translocon components SipB and SipC, which possess well-established effector-like activities. All fusions were introduced at their native genomic loci using λ Red recombineering^17^ to preserve native regulation and minimize artefacts associated with plasmid-based or inducible expression (**Supplementary Tables S1-2**).

### Differential effector expression under SPI-1– and SPI-2–inducing conditions

To assess expression and integrity of the HiBiT-tagged precursors, we first profiled bacterial lysates under conditions that induce either SPI-1- or SPI-2–associated effector production and secretion. HiBiT blotting of overnight (O/N) Luria–Bertani (LB) cultures, representing predominantly SPI-1-permissive conditions^18^, detected 26 of the 39 tagged effectors (67%) when analyzed across parallel gels (**Figs. 1-2** and **Supplementary Fig. S1A**). Growth in minimal low-phosphate, low-magnesium medium (LPM) under SPI-2–inducing conditions (pH 5.8)^19^ revealed 23 HiBiT-tagged effectors (59%) (**Figs. 1-2** and **Supplementary Fig. S1B**). Across these two conditions, eight effectors (GtgA, SipA-C, SopD, SopE, SopE2, and SteB) were detected exclusively under SPI-1-permissive conditions, corresponding largely to known SPI-1 effectors (**Fig. 1A, C**) (**Supplementary Table S1**). Conversely, five effectors (SteD, SifA, SpvC, SseK2, and SseI) were uniquely detected under SPI-2-inducing conditions. In total, 31 effectors (79%) were detectable on the initial gel set, yielding relative quantitative effector expression info (**Fig. 1 and Supplementary Fig. S1**).

**Figure 1.**
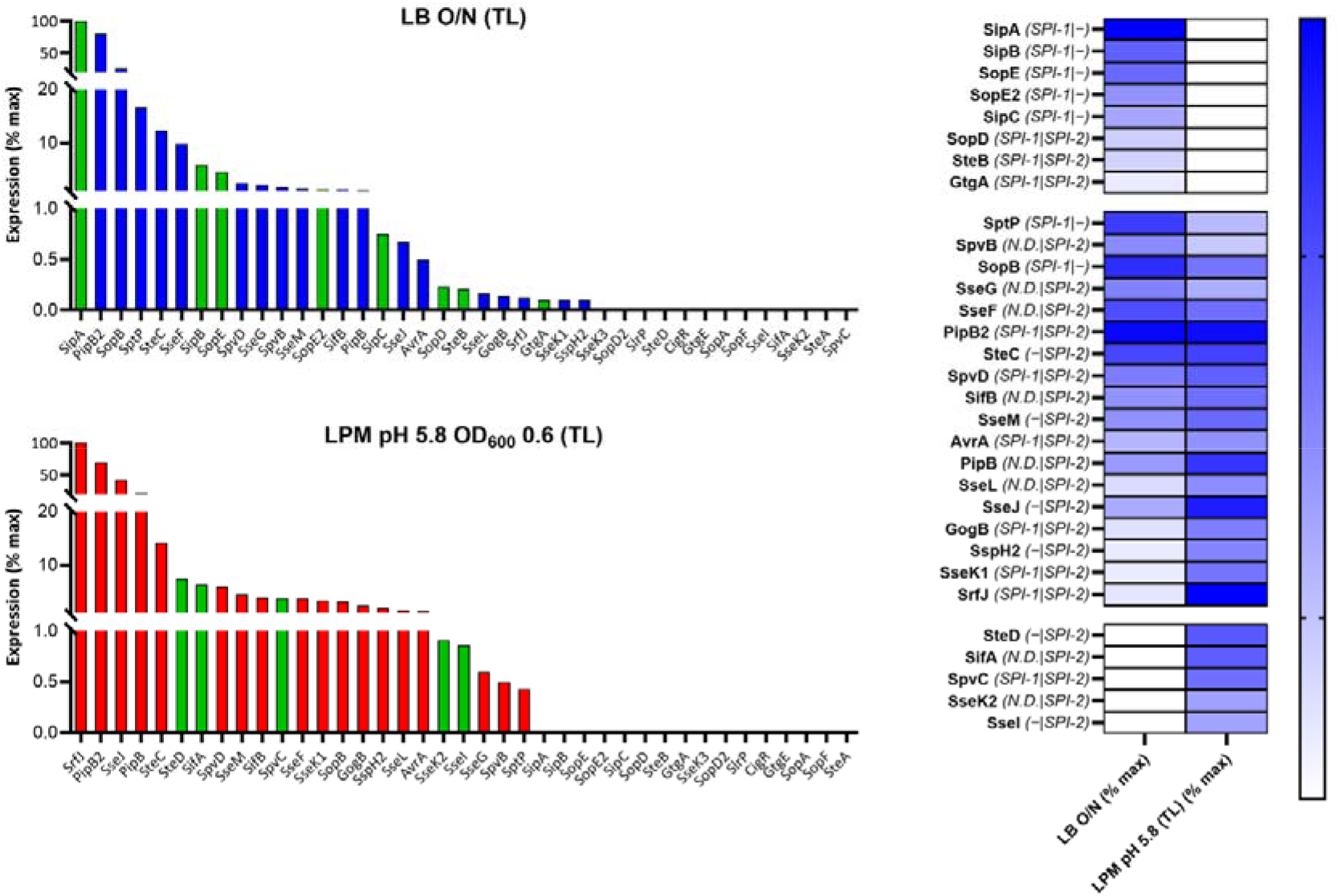
Expression profiling of endogenous HiBiT-tagged *Salmonella* type III effectors under SPI-1- and SPI-2-inducing conditions. HiBiT blotting data of whole-cell (total) lysates (TL) from *Salmonella enterica* serovar Typhimurium SL1344 strains encoding endogenous individual C-terminal HiBiT fusions of 39 type III effectors are shown in **Supplementary Figure S1**. HiBiT–LgBiT complementation was used for targeted detection of effector protein expression and/or secretion. **(A)** Relative effector expression levels during growth in LB overnight (O/N) cultures (SPI-1–permissive conditions). Bars represent HiBiT signal intensities from whole-cell lysates, normalised to the highest detected signal in this condition (% max). Green bars denote effectors detected exclusively under SPI-1–permissive growth; blue bars indicate effectors detected in both conditions. **(B)** Relative expression levels of the same HiBiT-tagged effector panel during growth in SPI-2-inducing, mildly acidic LPM medium (pH 5.8) to mid-log phase (OD_600_ = 0.6). Bars show HiBiT-normalised expression (% max) as in panel A. Green bars represent effectors detected exclusively in LPM medium (SPI-2–inducing conditions), whereas red bars indicate those detected in both conditions. **(C)** Heatmap summarising the expression profiles obtained from panels A and B. Effectors are grouped into three categories: (i) SPI-1-exclusive (detected only under SPI-1–permissive LB conditions) (8 effectors), (ii) detected in both conditions (18 effectors), and (iii) SPI-2-exclusive (detected only under SPI-2–inducing LPM conditions) (5 effectors). Color intensity represents log_2_(% max intensity × 100), providing a scaled, positive measure of relative expression, with white boxes indicating no detectable expression. The SPI-1/SPI-2 annotation beside each effector reflects its documented expression/secretion classification (SPI-1, SPI-2, dual SPI-1/SPI-2, or non-defined (N.D.))^2, 20^, corresponding to the categorisation provided in **Supplementary Table S1**.

**Figure 2.**
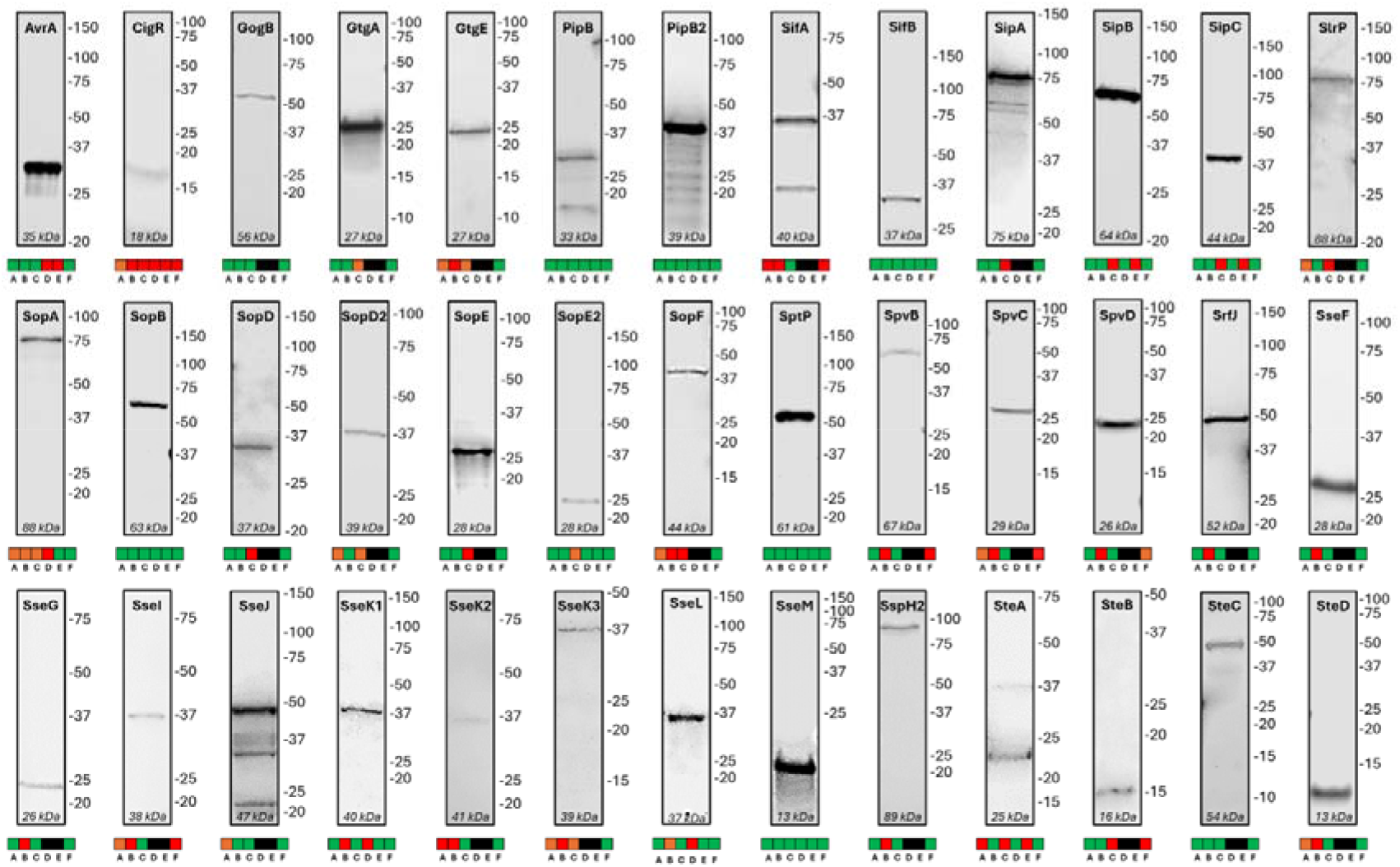
Comprehensive HiBiT-based survey of type III effector expression, secretion, and translocation. Representative HiBiT blots were obtained from three experimental conditions: (1) SPI-1–permissive LB conditions, including total lysates (TL) from early stationary phase cultures (OD_600_ = 2.0) or overnight cultures, along with the corresponding secreted fractions (secretomes); (2) SPI-2–inducing conditions using mildly acidic LPM medium (pH 5.8); and (3) infection-derived fractions from HeLa cells infected at a multiplicity of infection (MOI) of 50, separated into TX-100–insoluble pellets (representing bacteria-associated effectors) and TX-100– soluble supernatants (representing translocated effectors). For each effector, a representative blot from a condition in which a clear HiBiT signal was obtained is shown. The accompanying six-square panel indicates detection (green), detection at higher protein load (orange), no detection (red), or not determined (black) across: (A) LB TL (OD_600_ = 2.0 and/or LB O/N), (B) LB secretomes (O/N), (C) SPI-2–inducing conditions TL, (D) TX-100-insoluble pellet fraction (infected HeLa cells) TL, (E) TX-100-soluble supernatant (translocated effectors), and (F) real-time HiBiT translocation readout (infected HeLa cells). Representative blots obtained from LB OD_600_ = 2.0 TL: AvrA (PVDL0187), GtgA (PVDL0313), GtgE (PVDL0246), SipB (PVDL0183), SipC (PVDL0028), SlrP (PVDL0184), SopB (PVDL0024), SopD (PVDL0185), SptP (PVDL0034), and SteC (PVDL0263). Representative blots obtained from LB O/N TL: CigR (PVDL0045), SifB (PVDL0041), SipA (PVDL0328), SopD2 (PVDL0299), SopE (PVDL0267), SopE2 (PVDL0033), SopF (PVDL0278), SpvB (PVDL0425), SseF (PVDL0257), SseK1 (PVDL0248), SseL (PVDL0249), SseM (PVDL0046), and SteB (PVDL0311). Representative blots obtained from SPI-2–inducing conditions: GogB (PVDL0298), PipB (PVDL0251), PipB2 (PVDL0040), SifA (PVDL0312), SpvC (PVDL0426), SpvD (PVDL0427), SrfJ (PVDL0280), SseD (PVDL0270), SseG (PVDL0258), SseI (PVDL0259), SseJ (PVDL0275), SseK2 (PVDL0260), SseK3 (PVDL0289), SspH2 (PVDL0261), and SteD (PVDL0271). Representative blots obtained from additional conditions: SopA from infection-derived Triton X-100–insoluble pellet fractions from infected HeLa cells (PVDL0032; 8 h post-infection (hpi)) and SteA (PVDL0029; LB O/N secretome).

Increasing the protein load on additional gels and analyzing total lysates (TLs) from early stationary phase cultures, in addition to O/N LB cultures, together with the corresponding secretomes (see Methods), provided HiBiT blotting evidence for eight further effectors. In total, this yielded HiBiT-based expression validation for all 39 effectors, each detected under at least one of the tested growth conditions (**Figs. 1-2, Supplementary Fig. S1A-B**, and data not shown).

To characterise the secretion competence of the HiBiT-tagged effector strains, we first examined in vitro effector export under SPI-1–permissive conditions. LB O/N secretomes were analysed by HiBiT blotting alongside their corresponding total lysates to evaluate both effector expression and T3SS-1– dependent secretion capacity. Clear HiBiT-positive bands were detected in the LB O/N secretomes for a broad subset of 23 effectors (representing secretion of 70% of the 33 effectors detected in TLs under these conditions), including significant secretion of the canonical SPI-1 effectors SipA, SipB, SipC, SlrP, SopB, SopE, SopE2, and SptP, as well as several effectors with known dual SPI-1/-2 secretion capability (GogB, GtgA, PipB2, SteA, and SteB) (**Supplementary Table S1**). These findings are consistent with robust T3SS-1–mediated export in LB cultures (**Fig. 3**). Except for SteA and SlrP, all secreted effectors identified in the secretome were also detected in the corresponding total lysate samples. Under these conditions, sixteen effectors showed no detectable secretion in LB O/N secretomes. Notably, only a few of these exhibited (low) expression in total lysates, and all except SopF and SopA–both poorly expressed under all in vitro conditions tested–belong to the SPI-2-associated cohort.

**Figure 3.**
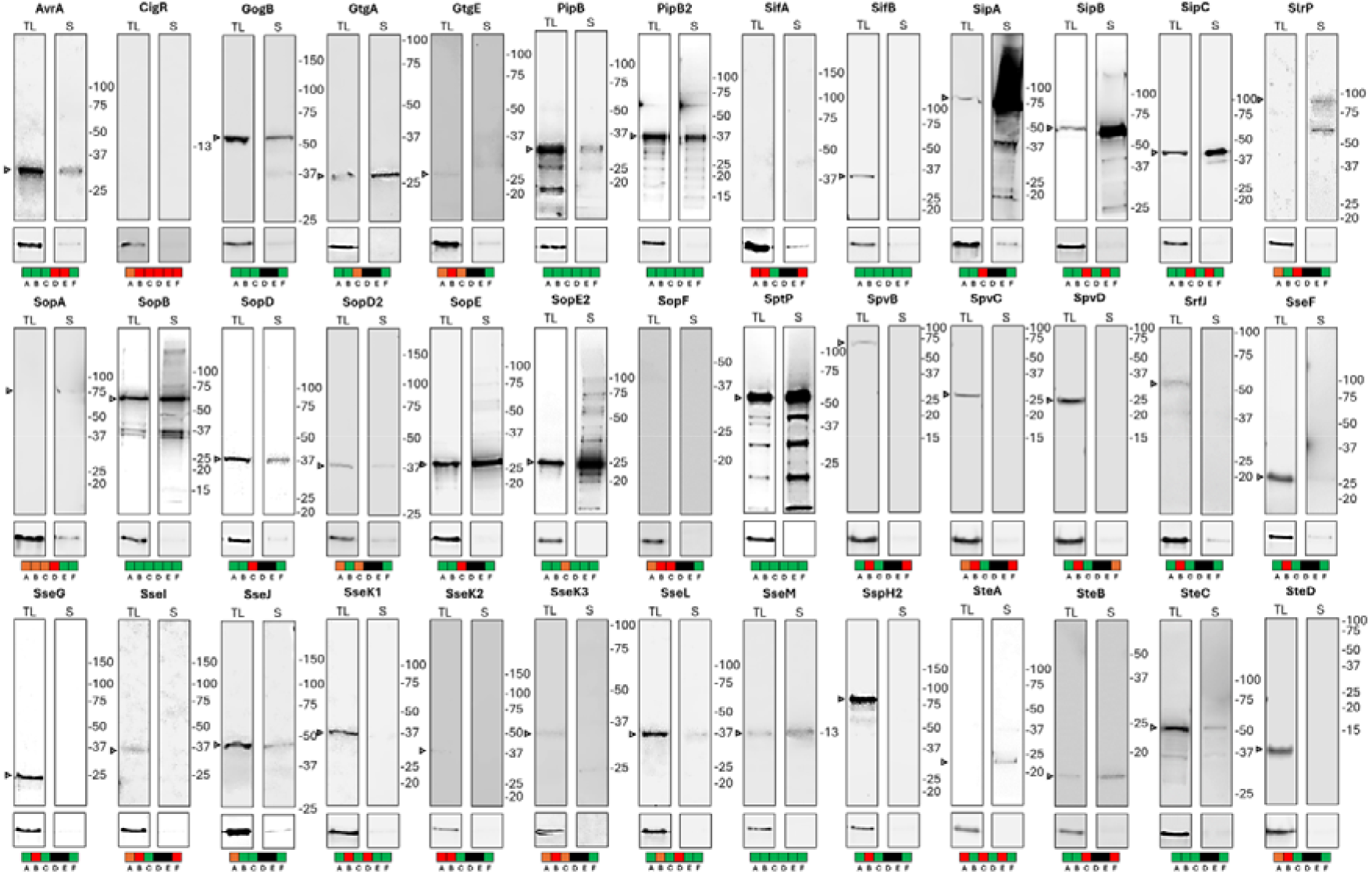
HiBiT blotting-based survey of type III effector secretion from overnight LB cultures. Representative HiBiT blots illustrating the secretion potential of all 39 *Salmonella* typedetected green fluorescent objects III effectors in overnight (O/N) LB cultures, which mainly supports SPI-1–dependent secretion. For each effector, total lysates (TL) are shown alongside their corresponding secretome (S) fractions, providing a comparative map of basal, SPI-1–mediated secretion capacity. TL samples are included to demonstrate expression relative to secretion under these conditions. As in **Fig. 2**, the accompanying six-square panel summarizes detection outcomes across the major experimental categories, with green indicating detection, orange detection at higher protein load, red no detection, and black not determined, across: (A) LB TL (OD_600_ = 2.0 and/or LB O/N), (B) LB secretomes (O/N), (C) SPI-2–inducing conditions TL, (D) TX-100-insoluble pellet fraction (infected HeLa cells) TLs, (E) TX-100-soluble supernatants (translocated effectors), and (F) real-time HiBiT translocation readout (infected HeLa cells). All HiBiT-tagged effectors were expressed from endogenously modified strains in the GFP-expressing SL1344 background (*hisG*46 P_tet_::*gfp*; PVD0002), except for SipA, SpvB, SpvC, and SpvD, which were expressed in the wild-type SL1344 background (PVD0001). Individual strain identifiers are listed in **Supplementary Table S2**.

Having established which effectors are secreted under SPI-1–permissive conditions, we next assessed the ability of C-terminally HiBiT-tagged effectors to translocate into host cells during infection. HeLa monolayers were infected with a selected panel of 10 representative effectors covering distinct SPI-associated secretion classes, and Triton X-100–soluble host cell fractions were analysed in parallel with matched bacteria-associated pellets recovered from the same culture samples. Five effectors—SopB and SptP (canonical SPI-1 effectors^2, 21, 22^), SseM^16^ and SifB^23^ (SPI-2-linked effectors^2, 20^), and PipB2^24^ (dual SPI-1/SPI-2 effector^2, 20^)—displayed clear HiBiT signals in the Triton X-100–soluble fraction, demonstrating (robust) host cell translocation under these infection conditions (**Fig. 4**). In contrast, SopA, SipC, and SopE2 (SPI-1 effectors), as well as SteA (dual SPI-1/SPI-2 effector^2, 14, 20^), were detectable only in the bacteria-associated fraction. CigR—previously reported as an SPI-2 substrate only in an *ssaL* mutant background, where SPI-2 secretion is deregulated^25^—did not yield detectable HiBiT signal in either fraction, indicating a lack of detectable expression and host cell translocation under these conditions (data not shown).

**Figure 4.**
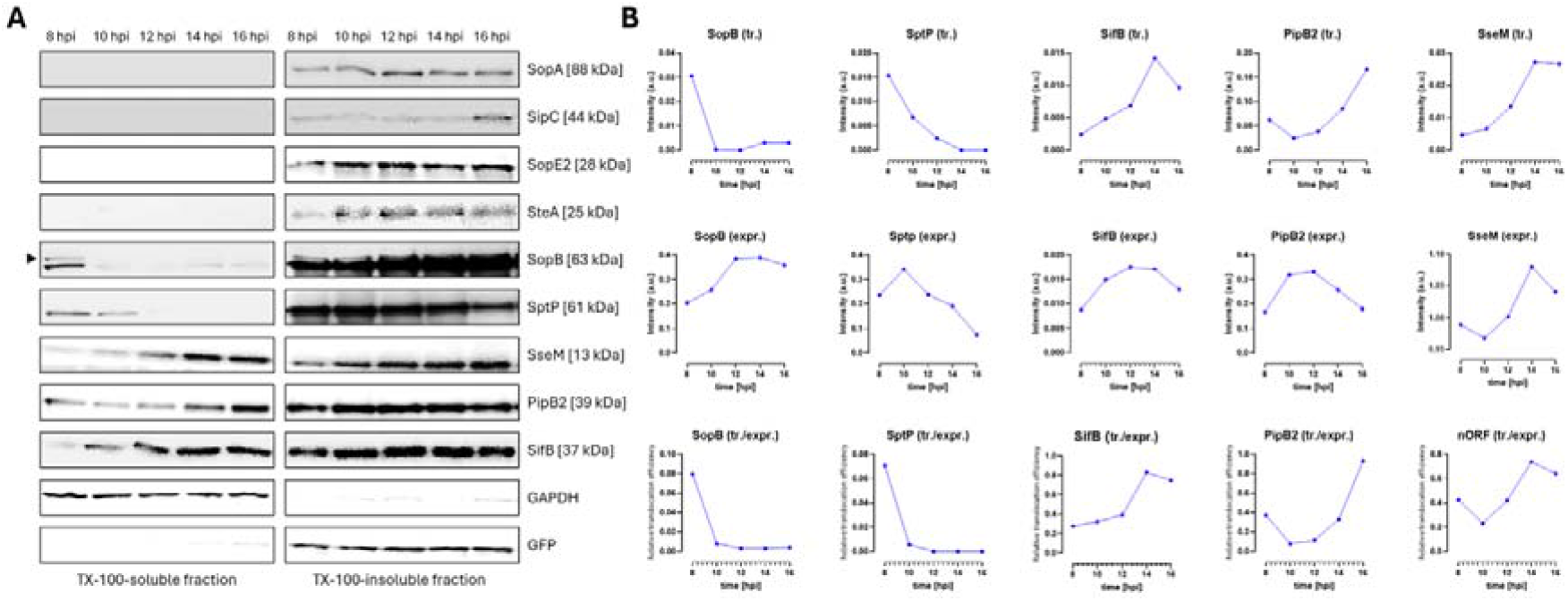
Kinetics of type III effector translocation into HeLa cells monitored by HiBiT blotting. HiBiT blotting analysis of 10 selected *Salmonella* type III effectors—SopA, SipC, SopE2, SteA, SopB, SptP, SseM, PipB2, SifB, and CigR—during infection of HeLa cells. CigR data are not shown due to the absence of a detectable HiBiT signal. **(A)** Representative HiBiT blots of Triton X-100–soluble (host cell cytosolic; translocated effectors) and Triton X-100–insoluble (bacteria-associated; expressed effectors) fractions collected at the indicated hours post-infection (hpi). GAPDH and GFP immunoblots are shown as representative and reproducible controls from the SopB experiment and were used as markers of the TX-100–soluble host cell fraction and as a normalization control for bacterial load, respectively. The black arrowhead denotes the mono-ubiquitinated form of the effector SopB, indicative of host-cell translocation. SopA, SipC, and SopE2 were detectable exclusively in the TX-100–insoluble fraction under the infection conditions tested (8–16 hpi), indicating bacterial expression but no detectable host cell translocation. **(B)** Densitometric quantification of HiBiT signals over time (background-corrected signal intensity; a.u.: arbitrary units). Upper panels depict translocated effector levels (tr.), quantified from the TX-100–soluble fraction. Middle panels show bacteria-associated effector levels (expr.), quantified from the TX-100–insoluble fraction. Lower graphs represent the relative translocation efficiency (tr./expr.), calculated as the ratio of translocated (TX-100–soluble) to bacteria-associated (TX-100–insoluble) HiBiT signal. Quantification is shown for those effectors displaying detectable host-cell translocation under the infection conditions tested (8–16 hpi) (SopB, SptP, SifB, PipB2, and SseM). HeLa cells were infected at an MOI of 50, and samples were processed as described in Methods. Abbreviations: tr., translocated effector (TX-100–soluble fraction); expr., bacteria-associated (expressed) effector (TX-100–insoluble fraction).

For the majority of effectors examined, HiBiT-positive bands were detected either in LB secretomes or in infection-derived host cell–associated fractions, consistent with efficient T3SS-dependent secretion and/or translocation of the tagged effector proteins. Although SopA was poorly detectable under the in vitro growth conditions tested, it became readily detectable in bacteria-associated fractions recovered after Triton X-100 lysis (**Figs. 2** and **4**), indicating that infection-associated cues promote its expression in the host environment.

Overall, all HiBiT-tagged effectors displayed detectable protein levels under at least one physiologically relevant growth or infection condition. Moreover, the observed secretion and translocation patterns were concordant with established SPI-1– and SPI-2–biased expression signatures and infection stage-specific effector deployment, with SPI-1–associated effectors predominating at earlier time points and SPI-2–associated effectors enriched at later stages of infection. Together, these endpoint infection analyses demonstrate that endogenous HiBiT-tagging is broadly compatible with effector synthesis and stability (as indicated by the preserved intact molecular weight profiles), and retains native SPI-1 and SPI-2 regulatory control as well as T3SS secretion and translocation competence, thereby providing a robust foundation for subsequent quantitative real-time profiling of effector translocation over an extended 24-hour infection course.

### Real-time profiling of infection dynamics during epithelial infection

To contextualize effector translocation kinetics within the broader infection landscape, we first assessed *Salmonella* proliferation dynamics in our previously characterized HeLa–SL1344 infection model^26^ using a constitutively eGFP-expressing (eGFP^+^) *S*. Typhimurium SL1344 strain^27^ at varying multiplicities of infection (MOIs) during the later stages of epithelial infection (**Fig. 5**).

**Figure 5.**
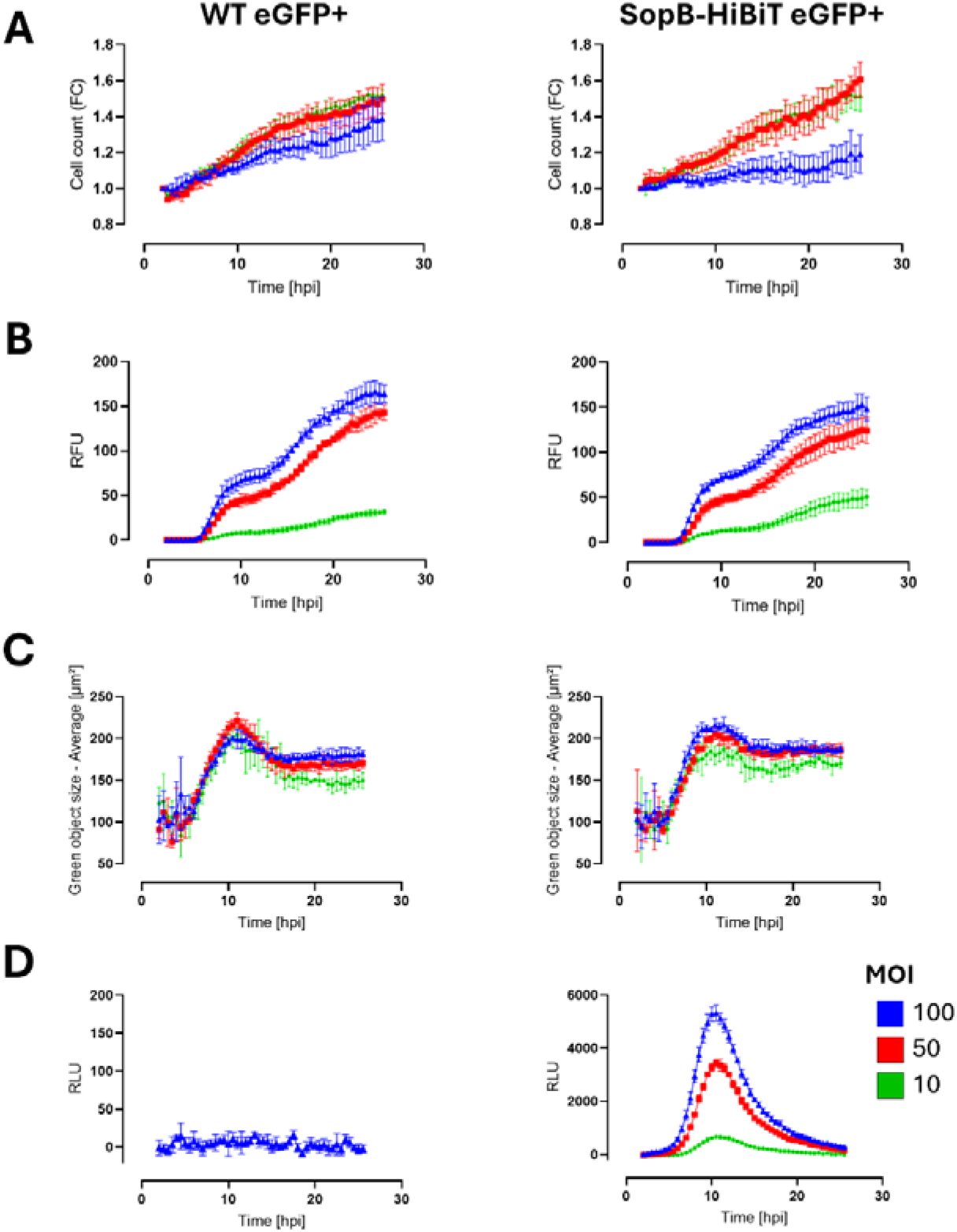
Characterization of the HeLa–*Salmonella* infection model for real-time monitoring of infection dynamics and effector translocation via NanoLuc complementation. HeLa cells stably expressing LgBiT were infected with a constitutively eGFP-expressing *Salmonella enterica* serovar Typhimurium SL1344 strain, either wild-type (WT eGFP+, left panels) or expressing chromosomally integrated *sopB-HiBiT* (SopB-HiBiT eGFP+, right panels), at multiplicities of infection (MOIs) of 10 (green), 50 (red), or 100 (blue). Infections were performed in quadruplicate, and real-time acquisition was conducted for 24 h using a multimode plate reader under environmental control. Data acquisition started 2 h post-infection (hpi). Top-to-bottom panels show: (i) fold change in HeLa cell count based on label-free automated brightfield imaging (10x objective); (ii) green fluorescence (RFU) as a proxy of bacterial load; (iii) average size (µm^2^) of detected green fluorescent objects; and (iv) luminescence signal (RLU) either background levels or SopB-HiBiT translocation into LgBiT-expressing HeLa cells via NanoLuc complementation. Vivazine Live Cell Substrate was added following the gentamicin protection step and prior to acquisition to enable continuous luminescence detection throughout the infection course, starting at 2 hpi. Data points (every 30’) represent the mean ± SD of quadruplicates. WT-infected controls (left) confirm minimal background luminescence in the absence of HiBiT-tagged effectors.

Real-time multimodal imaging under controlled environmental conditions enabled simultaneous acquisition of fluorescence, luminescence, and bright-field images, allowing bacterial replication to be monitored via total green fluorescence (RFU) and object counts (**Fig. 5B**), while host cell status was monitored by automated confluence and cell-count analysis (see Methods and **Fig. 5A**).

Using these complementary readouts, we observed a characteristic biphasic progression in intracellular bacterial load (**Fig. 5B**). A sharp increase in bacterial signal was detected between 5 and 9 hours post-infection (hpi), corresponding to the onset and peak of cytosolic hyper-replication following escape from the *Salmonella*-containing vacuole (SCV)^28, 29, 30^. This observation is consistent with previous studies in cultured epithelial cells reporting that by ∼8 hpi, in approximately 5–20% of infected epithelial cells, the SCV membrane becomes compromised, allowing bacteria to escape into the nutrient-rich cytosol, where they undergo rapid hyper-replication with doubling times of ∼20 minutes and account for up to half of the total intracellular bacterial population^28, 29, 31^. This cytosolic subpopulation is not only spatially distinct but also exhibits a distinct virulence program, characterized by re-expression of flagella and reactivation of T3SS-1, a phenomenon referred to as the “second wave” of SPI-1 activity^13, 29, 30^. Consistent with this, the *Salmonella* T3SS-1 apparatus has been shown to be required for early vacuolar lysis and efficient colonization of the epithelial cytosol, whereas deletion of a suite of individual SPI-1 effectors (SptP, SopE, SopE2, SopB, AvrA, SopA, SipA) does not significantly impair vacuolar escape, implying the involvement of T3SS-1 itself and additional mechanisms in this process^29^. These cytosolic bacteria are therefore expected to contribute substantially to T3SS-1–dependent effector delivery during the post-invasion phase of infection, whereas bacteria remaining within intact vacuoles predominantly engage T3SS-2– mediated effector translocation during later stages.

Accordingly, the pronounced increase in bacterial signal reflects rapid intracellular expansion within a subpopulation of infected cells that accounts for the majority of net bacterial replication. This initial amplification phase was followed by a plateau from ∼10–15 hpi, coinciding with the onset of host cell rounding, cytotoxicity, and lysis-associated loss of cytosolic bacteria. As gentamicin prevents reinfection of neighboring cells, this plateau likely reflects a dynamic balance between cytosolic bacterial loss and the slower, SPI-2–dependent replication of bacteria that remain within intact vacuoles. A second expansion phase became apparent from approximately 15 hpi onwards, consistent with continued replication of the vacuolar population, with indications of stabilization toward the end of the 26-hour infection time course (**Fig. 5B**).

Object size analysis supported this interpretation: average GFP-positive object area peaked at ∼11 hpi and subsequently decreased by ∼10–20% (**Fig. 5C**), consistent with host cell rounding, reduced projected cell area, and the transition from cytosolic hyper-replication to a state in which the remaining bacteria are predominantly contained within intact vacuoles. Together, these data highlight three main operational phases of intracellular infection dynamics in epithelial cells: (i) cytosolic hyper-replication, (ii) a transition/plateau phase driven by host cell damage and progressive loss of cytosolic bacteria, and (iii) sustained vacuolar replication. These distinctions are essential for interpreting the kinetic waves of effector translocation observed in the real-time assays.

Further supporting this progression, infection dynamics were also profiled using constitutively eGFP-expressing SL1344 mutants lacking key virulence functions (**Supplementary Fig. S2**): a Δ*invA* (T3SS-1-deficient), Δ*ssaV* (T3SS-2-deficient), and *sifA* translational (triple) knockout (tko, Δ*sifA*) strain (**Supplementary Table S3**). As expected, the Δ*invA* strain was impaired in epithelial invasion and exhibited only background fluorescence throughout the time course, consistent with its inability to initiate intracellular infection via T3SS-1-mediated entry (**Supplementary Fig. S2B**). In contrast, Δ*ssaV* mutants displayed a pronounced early cytosolic amplification phase but failed to undergo the later secondary expansion observed in wild-type infections (**Supplementary Fig. S2C**). This phenotype is consistent with previous reports showing that the *Salmonella* T3SS-2, while essential for SCV biogenesis and maintenance, is dispensable for cytosolic replication, and that bacteria lacking the T3SS-2 exhibit a delayed replication defect in epithelial cells due to their inability to establish a stable vacuolar niche^28, 32^. Accordingly, the absence of a second replication phase in Δ*ssaV* infections confirms the requirement for T3SS-2–dependent vacuolar remodeling for sustained intracellular replication. Notably, the Δ*sifA* (tko) strain exhibited a more continuous increase in GFP signal and consistently higher overall fluorescence at comparable MOIs (**Supplementary Fig. S2D**). The increased GFP signal observed reflects a net increase in Δ*sifA* bacterial replication driven by a higher proportion of infected cells harbouring hyper-replicating bacteria. This pattern is indicative of premature SCV rupture, early cytosolic dominance, and enhanced cytosolic hyper-replication, consistent with loss of SifA-mediated SCV membrane integrity and the well-established cytosolic phenotype of SifA-deficient strains in epithelial cells^33, 34^. Together, these findings validate the utility of our real-time imaging platform for dissecting infection phase-specific dynamics and demonstrate that characteristic replication signatures can be leveraged to functionally profile *Salmonella* virulence mutants.

### Real-time profiling of effector translocation via NanoLuc complementation

To monitor T3SS-mediated effector translocation in real-time during the later stages of epithelial infection, we used HeLa cells stably expressing the LgBiT fragment of NanoLuc ^11, 35^. These cells were infected with the panel of *S*. Typhimurium SL1344 strains encoding endogenously HiBiT-tagged effectors. Real-time multimodal imaging of infections with the library revealed infection dynamics that were largely comparable to those of the wild-type (WT) eGFP-expressing SL1344 strain, as assessed by total green fluorescence (RFU) and green object counts (**Supplementary Fig. S3**). These findings indicate that endogenous C-terminal HiBiT-tagging generally preserves native infection dynamics and does not markedly interfere with bacterial invasion or intracellular proliferation. Deviations from wild-type behaviour were, however, apparent in strains carrying HiBiT-tagged versions of the SPI-1 translocon components SipB and SipC. Whereas SipC-HiBiT infections were associated with a reduced intracellular bacterial load, reflected by a lower overall GFP signal, SipB-HiBiT infections exhibited a shortened cytosolic hyper-replication phase followed by a prolonged plateau. These changes occurred despite SipB infection profiles otherwise resembling those of canonical SPI-1 effectors, including similar object size dynamics and comparable effector translocation kinetics as assessed by luminescence, as discussed below.

Given that SipB, together with SipC, forms the translocon pore required for sustained T3SS-1– mediated effector delivery, even subtle perturbations introduced by C-terminal tagging may influence the invasion efficiency or persistence of post-invasion T3SS-1 activity. Previous studies have shown that tagging of translocon proteins, including SipB and SipC, can be variably tolerated depending on tag identity, size, and placement, and may, in some contexts, interfere with translocon function, necessitating alternative tagging strategies^36, 37^. Such effects are expected to primarily impact the size or stability of the cytosolic bacterial subpopulation, resulting in altered population-level replication dynamics without grossly affecting overall effector translocation profiles. With infection dynamics largely preserved, the NanoLuc complementation readout was used to directly quantify effector translocation in real-time.

Upon successful translocation into the host cytosol via T3SS-1 or T3SS-2, the HiBiT-tagged effector binds the host-expressed LgBiT fragment with high affinity (*K_D_* ∼0.7 nM), reconstituting a functional NanoLuc luciferase. In the presence of a furimazine-based substrate, this enzyme catalyzes substrate oxidation, producing a bright luminescent signal proportional to the amount of translocated effector^11, 35^. As previously demonstrated for SipA-HiBiT^11^, this strategy ensures that signal generation is restricted to the host cytosol. While this system has been successfully applied to detect effector translocation at early infection stages, its suitability for continuous, long-term profiling, as well as for profiling at later stages of infection, had not yet been explored. To assess both the feasibility and sensitivity of real-time effector translocation monitoring for extended-duration measurements and for profiling at later stages of infection, we first evaluated the use of Nano-Glo Vivazine, a pro-substrate that enables sustained signal generation through gradual hydrolysis in the culture medium, thereby maintaining a consistent supply of active substrate.

We initially validated this approach by profiling SopB, a canonical SPI-1 effector (**Fig. 5**). Notwithstanding a signal-to-noise ratio >50 at 2 hpi, luminescence peaked only around 9 hpi before declining, indicative of a prolonged phase of SopB translocation (**Fig. 5**). As hyper-replicating cytosolic *Salmonella* re-express flagella and the T3SS-1 at later stages of epithelial infection, this profile is consistent with the second wave of sustained T3SS-1–mediated SopB delivery^13^. This timing also aligns with endpoint HiBiT blotting of Triton X-100–soluble host cell fractions, which still showed evidence of translocated SopB at 8 hpi (**Fig. 4**), demonstrating concordance between real-time luminescence measurements and biochemical endpoint detection. Together, these results establish the suitability of NanoLuc complementation using vivazine for sensitively tracking effector injection kinetics over extended infection periods and provide a reference framework for interpreting translocation dynamics across effectors.

Building on this validation, we conducted live-cell monitoring from 2 to 26 hpi for all HiBiT-tagged effectors in their eGFP^+^ WT, Δ*invA*, and Δ*ssaV* mutant backgrounds (**Fig. 6** and **Supplementary Fig. S4**). Δ*invA* strains served as a negative control for effector injection and displayed luminescence levels at or near background. The resulting luminescence profiles across the effector library revealed distinct translocation patterns, enabling quantitative classification of SPI-1–restricted, dual-system, and SPI-2–specific effector translocation behaviors. SPI dependence classification was based on statistically significant reductions in area-under-the-curve (AUC) values in WT compared to Δ*invA* and Δ*ssaV* backgrounds across biological replicates (**Supplementary Table S5**).

**Figure 6.**
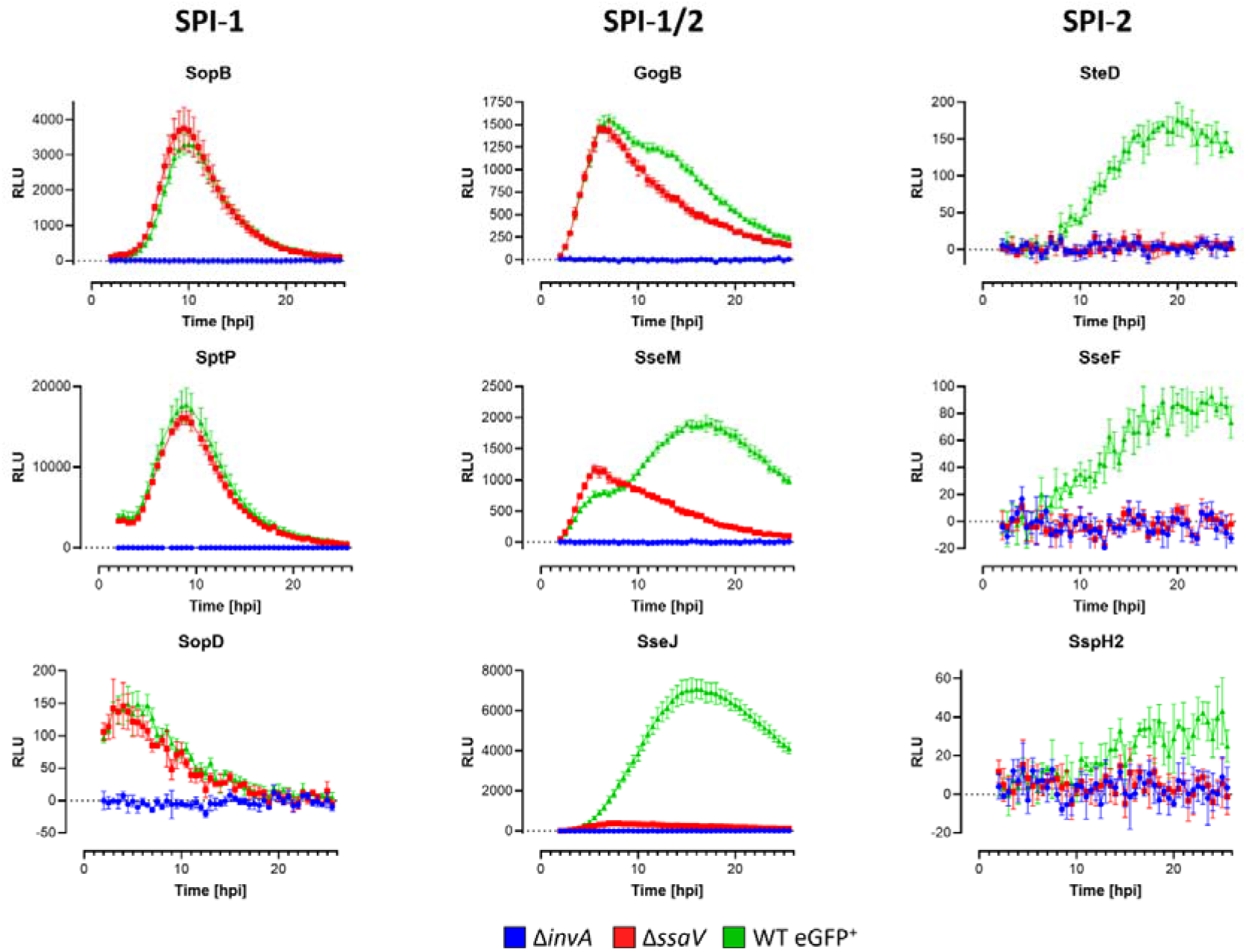
Real-time NanoLuc-based profiling of effector translocation dynamics in wild-type and secretion system-deficient backgrounds during epithelial infection. HeLa cells stably expressing LgBiT were infected with endogenously HiBiT-tagged *Salmonella enterica* serovar Typhimurium SL1344 effector strains in three genetic backgrounds: wild-type (WT) SL1344 carrying constitutive eGFP^+^ (*hisG*46 P_tet_ ::gfp; PVD0002; green), the T3SS-1–deficient *ΔinvA::cat* mutant (PVD0316; blue), and the T3SS-2–deficient *ΔssaV::cat* mutant (PVD0317; red). Luminescence profiles (RLU), reflecting NanoLuc complementation following effector translocation into the host cytosol, were recorded from 2 to 26 hours post-infection (hpi). Panels show a representative subset of effectors grouped by secretion system dependence—SPI-1 (SopB, SptP, SopD; left), SPI-1/2 (GogB [25% SPI-2 contribution], SseM [56%], SseJ [95%]; middle), and SPI-2 (SteD, SseF, SspH2; right)—as determined by phase-resolved AUC analysis (see Methods and **Supplementary Table S5**). The complete set of translocation profiles for all 39 effectors is shown in **Supplementary Figure S4**. Infections were performed at a multiplicity of infection (MOI) of 50, and real-time luminescence monitoring was conducted using a Spark Cyto 400 multimodal plate reader under environmental control in the presence of Nano-Glo Vivazine. Luminescence values were background-subtracted (non-infected HeLa). Data points represent mean ± SD of technical quadruplicates. Individual strain identifiers and effector annotations are provided in Supplementary Table S2.

Overall, 32 of the 39 effectors examined produced luminescence signals reproducibly above background in the WT (GFP^+^) background, confirming their successful translocation into HeLa cells under native expression conditions (**Supplementary Figs. S3 and S4, Supplementary Table S5**). In contrast, seven HiBiT-tagged effectors—CigR, SifA, SpvB, SpvC, SpvD, SseI, and SteB—did not reach the predefined AUC threshold for detectable translocation (see Methods and **Supplementary Table S5**) and were therefore classified as not detectably translocated in this epithelial cell model. Although several effectors—most notably the SPI-2 effectors SseG, SseK2, and SspH2—exhibited relatively modest signal amplitudes during long-term vivazine-based monitoring in the WT background (e.g., signal-to-noise ratios (S/N) at 24 hpi of 20 for SseK2, 34 for SseG, and 43 for SspH2), their translocation was independently confirmed using short-term, high-sensitivity real-time assays with furimazine at 24 hpi. These furimazine-based measurements yielded markedly higher S/N values (e.g., >360 for SseK2, 1030 for SseG, and 713 for SspH2) and demonstrated statistically significant translocation levels in the WT compared to the Δ*invA* background (adjusted P < 0.01, paired t-test), consistent with the higher peak sensitivity of furimazine despite its limited temporal stability (**Supplementary Figure S5**). These results demonstrate strong concordance between the long-term vivazine-based and short-term furimazine-based translocation assays. In contrast, no significant translocation signal was detected at 24 hpi for the SPI-2 effector SseI or the dual SPI effector SteB using either detection method. Conversely, SteC, which showed the highest vivazine-derived luminescence at 24 hpi among the tested effectors, also produced the strongest signal in the furimazine assay (adjusted P < 0.001), further supporting consistency between the two detection approaches (**Supplementary Fig. S5**).

### Phase-resolved classification of effector translocation dependence on SPI-1 and SPI-2

To determine secretion system dependence and quantify the relative contribution of each T3SS to effector delivery, translocation profiles were compared between the WT, Δ*invA*, and Δ*ssaV* strain backgrounds across mid-stage (2–10 hpi) and late-stage (10–24 hpi) infection. This phase-resolved real-time framework enabled refinement of effector classifications that were previously undetermined or inconsistently defined in the literature and resolved three principal dependency classes, revealing extensive functional overlap between SPI-1 and SPI-2 (**Supplementary Table S5**). Based on these phase-specific comparisons, we identified 21 effectors with partial or exclusive SPI-2 dependency, including seven effectors—PipB, SseF, SseG, SseK2, SseK3, SspH2, and SteD—that showed strictly T3SS-2-dependent translocation under these conditions. This confirms and extends previous reports for PipB, SseF, SseG, and members of the SseK family (SseK2/3) and identifies these effectors as late-stage cargo whose peak WT translocation windows (defined as ≥90% of maximal signal) occur exclusively during late-stage infection (≥13 hpi) and persist through the remainder of the infection course (**Fig. 6** and **Supplementary Table S5**).

Conversely, ten effectors were classified as exclusively T3SS-1 dependent (GtgE, SipA, SipB, SipC, SopA, SopB, SopE2, SopF, SptP, and SrfJ), showing significant WT translocation during the mid-stage infection phase with no statistically supported SPI-2 contribution during late-stage infection. This classification is largely consistent with previous reports for canonical invasion-associated effectors (**Supplementary Tables S1** and **S5**). In contrast, GtgE—previously reported as a dual-T3SS substrate (**Supplementary Tables S1** and **S5**)—was here classified as SPI-2-independent, reflecting undetectable SPI-2 translocation levels observed for this effector in the epithelial infection model used. SopE, whose translocation has been reported to depend exclusively on SPI-1, was likewise classified as a dual-system effector in this analysis, although its relative SPI-2 contribution remained low (∼14% of total WT translocation). Because SPI-2 contribution was inferred from differences between WT and Δ*ssaV* translocation levels, variability in infection efficiency or intracellular bacterial replication between strain backgrounds may influence quantitative estimates for low-amplitude signals, particularly for effectors with minor late-stage translocation.

Interestingly, fifteen effectors were classified as dual SPI-1/SPI-2-dependent (AvrA, GogB, GtgA, PipB2, SifB, SlrP, SopD, SopD2, SopE, SseJ, SseK1, SseL, SseM, SteA, and SteC), corresponding to 38% of the annotated effector repertoire (15/39) and 47% of the detectably translocated effectors (15/32), a substantially higher proportion than generally reported in the literature (**Supplementary Tables S1** and **S5**). Of these, only eight have previously been demonstrated to exhibit intrinsic dual-system translocation capacity, whereas the remaining effectors were primarily classified as SPI-2-associated. Notably, several effectors previously considered strictly SPI-2-dependent—SifB, SopD2, SseJ, SseL, SseM, and SteC—showed statistically supported T3SS-1-dependent delivery, with significant translocation detectable in the Δ*ssaV* background during mid-stage infection (2–10 hpi), consistent with contribution from the second wave of T3SS-1 activity.

### Hierarchical organization of effector delivery based on peak translocation timing

To further resolve temporal patterns within these dependency classes, we next ordered effectors according to the timing of their maximal translocation (T_max_; **Supplementary Table S5**) and visualized their normalized translocation profiles as matrix plots (**Fig. 7** and **Supplementary Fig. S6**). In the WT background, effectors segregated into early-, intermediate-, and late-delivered groups, revealing a hierarchical organization of effector deployment across the infection course (2-24 hpi). SPI-1– dependent effectors reached maximal translocation significantly earlier (median T_max_ = 9 hpi), whereas SPI-2–dependent effectors peaked predominantly during late-stage infection (median T_max_ = 22.5 hpi), with dual-system substrates displaying intermediate kinetics (median T_max_ = 16 hpi). In the absence of T3SS-2 (Δ*ssaV*), late-stage peaks were selectively lost for SPI-2–dependent and dual-system effectors, whereas SPI-1–restricted substrates retained their characteristic WT translocation profiles (**Supplementary Fig. S6**). Notably, among dual-system effectors, the SPI-1–associated T_max_ inferred from Δ*ssaV* profiles occurred earlier than for SPI-1–restricted effectors (median 6.5 hpi versus 9.5 hpi, respectively), suggesting that dual substrates may be preferentially deployed during early intracellular SPI-1 activity, including the post-invasion and cytosolic ‘second wave’ phases. Together, these data demonstrate that effector delivery follows a structured temporal hierarchy reflecting both secretion system usage and effector-specific regulation, rather than a simple binary switch between SPI-1 and SPI-2 activity.

**Figure 7.**
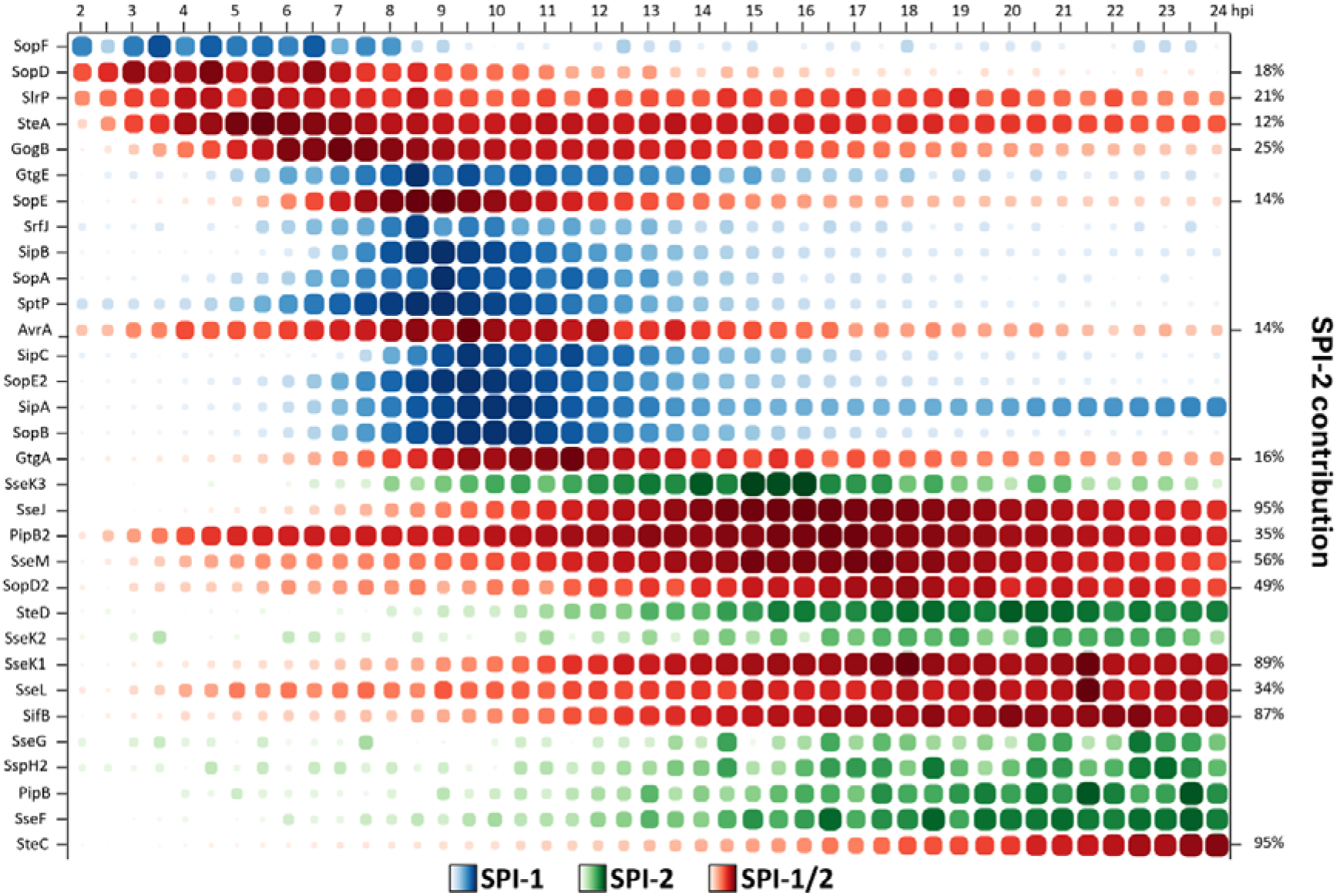
Hierarchical organization of effector translocation timing based on peak delivery. Matrix plots depict the temporal progression of effector translocation in a wild-type background, based on real-time HiBiT luminescence measurements normalized to the maximal signal observed for each effector (% max). Values represent means of technical quadruplicates. Effectors are ordered vertically according to ascending time of maximal translocation (T_max_), thereby revealing a hierarchy of early-, intermediate-, and late-delivered substrates. Only effectors that met the criteria for detectable translocation are shown (32 effectors). Each column represents a time point from 2 to 24 hours post-infection (hpi). Dot size and color intensity indicate the relative proportion of maximal translocation at each time point (% max). Effectors are color-coded by secretion system dependence as determined by phase-resolved AUC analysis: SPI-1–dependent (blue), SPI-2– dependent (green), and dual SPI-1/SPI-2–dependent (red). For dual-system substrates, the relative contribution of SPI-2 to total cumulative translocation (calculated from WT vs. ΔssaV AUC values) is indicated on the right-hand axis as the percentage of SPI-2–dependent delivery. Across the WT dataset, median T_max_ values were 9 hpi for SPI-1–dependent effectors, 16 hpi for dual-system effectors, and 22.5 hpi for SPI-2– dependent effectors, quantitatively reflecting the temporal stratification of secretion system usage.

### System-wide contribution of individual effectors to cumulative translocation output

To integrate the classifications at the level of the entire effector repertoire, we next quantified the relative contribution of each effector to total cumulative translocation over the full infection course (2–24 hpi) (Fig. 8 and Supplementary Table S5).

**Figure 8.**
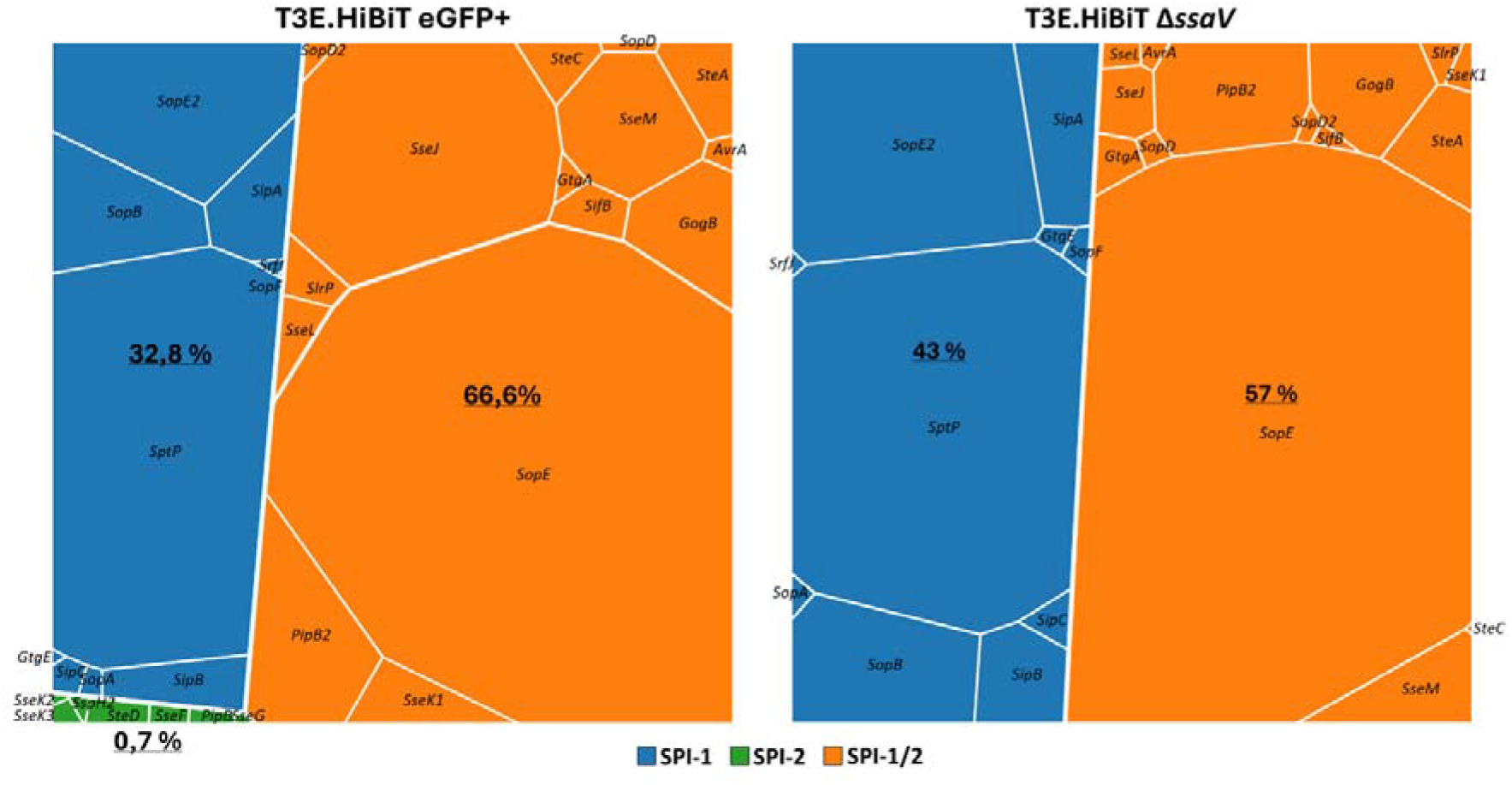
System-wide contribution of SPI-1, SPI-2, and dual-system effectors to cumulative type III effector delivery in wild-type (WT) and Δ*ssaV* backgrounds. Voronoi treemaps depict the relative contribution of individual effectors to total effector translocation, quantified as mean luminescence area-under-the-curve (AUC) from 2–24 hours post-infection (hpi) (**Supplementary Table S5**). Polygon area is proportional to cumulative WT (left) or ΔssaV (right) AUC for each effector. Polygons are colored according to secretion system dependence: SPI-1–dependent (blue), SPI-2–dependent (green), or dual SPI-1/SPI-2–dependent (orange), as determined by phase-resolved statistical comparisons (see Methods). In the WT background, total effector delivery is dominated by dual-system substrates (66.6%), with SPI-1–restricted effectors contributing 32.8% and SPI-2–restricted effectors contributing less than 1% of the total signal. In the absence of SPI-2 (ΔssaV), SPI-2–restricted effector translocation is abolished, and the cumulative contribution of dual-system substrates is reduced, reflecting loss of their SPI-2–mediated delivery component. Percentages indicate the fraction of total cumulative AUC contributed by each dependency class.

In the WT background, cumulative effector delivery was dominated by effectors classified as dual SPI-1/SPI-2 substrates, which together accounted for approximately two-thirds of the total translocation signal, whereas strictly SPI-1–dependent effectors contributed roughly one-third and strictly SPI-2–dependent effectors contributed less than 1% of total cumulative AUC (**Fig. 8A** and **Supplementary Table S5**). In the Δ*ssaV* background, SPI-2–restricted translocation was abolished, and cumulative delivery by dual-system substrates was substantially reduced (by ∼10% of total WT signal), reflecting loss of their SPI-2–mediated component, with a corresponding increase in the relative contribution of SPI-1–dependent substrates (**Fig. 8**). These data indicate that SPI-2 activity contributes to the total effector load throughout infection, not only through exclusive SPI-2 substrates but also substantially via a large cohort of dual-system effectors. These distributions highlight that while SPI-2 effectors are essential for vacuolar maintenance and intracellular persistence, they constitute only a minor fraction of the total effector mass delivered, whereas SPI-1 and dual substrates dominate the cumulative translocation output.

The ten effectors identified as exclusive T3SS-1 substrates are largely involved in the initial events of host cell manipulation. SopB, a phosphoinositide phosphatase, is known to activate the pro-survival kinase Akt to prevent apoptosis in infected cells, a function that has been shown to persist beyond invasion^38^. The real-time data further indicate that *Salmonella* continues to deliver SopB well after invasion has concluded, with WT translocation levels reaching near maximal levels toward the end of mid-stage infection, with a peak translocation window centered around 9-11 hpi.

The seven effectors that exclusively depend on the T3SS-2 constitute the canonical intracellular effector repertoire of *Salmonella*. Their translocation is minimal during mid-stage infection (2–10 hpi), accounting for only 8.4% of total T3SS-2–dependent translocation, but increases sharply during late-stage infection (≥10 hpi), when they account for 91.6%. WT translocation maxima for these effectors are broadly distributed between 16 and 23 hpi and are confined to the late infection phase for all seven effectors, consistent with their classification as strictly SPI-2 dependent (**Supplementary Table S5**).

Effectors such as SseF and SseG are physically and functionally linked and act in concert with host factors to tether the SCV to the Golgi apparatus^39^. Their late-stage translocation kinetics are consistent with this role in juxtanuclear positioning of the SCV and in *Salmonella*-induced filament (Sif) biogenesis^20, 40^. Similarly, members of the SseK family of arginine glycosyltransferases display predominantly late-stage translocation, with SseK3 targeting host Rab GTPases to the Golgi to suppress cytokine secretion^41^.

The *spv* locus (*spvB, spvC*, and *spvD*) has been linked to systemic virulence, and *spv* genes contribute to intracellular survival and replication in macrophages^42^. These effectors are translocated primarily via the SPI-2–encoded T3SS, and their expression and delivery are optimized for the intracellular environment of professional phagocytes^43^. In the HeLa epithelial infection model used here, environmental cues required for robust induction of the *spv* operon are likely insufficient, resulting in translocation levels below the standard detection threshold. Consistent with this interpretation, weak but statistically supported T3SS-2–dependent translocation of SpvD became detectable when lower AUC cutoffs were applied, indicating that Spv effector delivery can occur in epithelial cells, albeit at levels substantially lower than those typically observed in macrophage infection models. The identification of effectors that are translocated by both the T3SS-1 and T3SS-2 represents a significant shift in understanding the “secretion dichotomy”. This versatility allows *Salmonella* to utilize the same protein at different stages of infection, likely targeting different host pathways or localizations^44^.

SteA is one of the few effectors historically recognized as a substrate for both systems^14^. Our real-time data confirm its high-versatility profile, with significant translocation observed throughout the 24-hour course. SteA’s N-terminal 10 amino acids contain a signal sufficient for recognition by both the SPI-1 and SPI-2 secretion machineries^14^. Accordingly, early T3SS-1-mediated delivery coincides with the invasive phase of infection, whereas sustained T3SS-2-mediated translocation during late-stage infection contributes to SCV membrane dynamics and phenotypes associated with vacuolar remodeling^45^.

PipB2 exhibits a similar dual-dependency^46^. Although originally described as an SPI-2 effector^24^ and recognized as a key factor in kinesin-1 recruitment to the SCV^47^, real-time translocation profiling revealed substantial T3SS-1–mediated translocation during early and mid-stage infection, corresponding to the post-invasion and the second wave of SPI-1 activity. This T3SS-1–mediated delivery likely ensures that the molecular machinery required for kinesin recruitment and membrane tubulation is in place as soon as the SCV forms, rather than being delayed until full induction of the SPI-2 secretion system.

Previous studies reported that PipB2 translocation in epithelial cells is predominantly SPI-1-dependent, whereas in macrophages it occurs via SPI-1 during invasion and via SPI-2 at later time points^46^. In contrast, our longitudinal profiling in epithelial cells reveals clear SPI-2–dependent delivery during late-stage infection, closely resembling the biphasic translocation pattern previously described in macrophages. In line with SteA, the N-terminal 10 amino acids of PipB2 are sufficient for translocation by both T3SS-1 and T3SS-2^46^, supporting an intrinsic dual-recognition capacity that likely underlies the biphasic secretion profiles observed across the infection course.

The classification of SifB, SopD2, SseJ, SseL, SseM, and SteC as dually translocated effectors represents a refinement of current effector annotations enabled by this real-time profiling approach (**Supplementary Table S5**). Several of these effectors illustrate how dual-system translocation can expand functional activity across infection stages.

SopD2, previously characterized as an SPI-2 substrate required for Sif formation and intracellular replication in macrophages^48^, also showed statistically supported T3SS-1–dependent translocation during the mid-stage of epithelial infection in our dataset. This finding suggests that SopD2-mediated modulation of host endocytic trafficking may be initiated earlier during infection than previously appreciated.

SseL, a deubiquitinase implicated in suppression of NF-κB–dependent inflammatory signaling^49^, likewise displayed significant T3SS-1–dependent delivery during mid-stage infection. Early translocation of SseL may therefore contribute to dampening host inflammatory responses shortly after invasion, potentially complementing the activity of SPI-1 anti-inflammatory effectors such as AvrA^50^.

SseM, a recently described SPI-2 effector co-transcribed with *pipB2*, targets the host dystrophin-associated protein complex (DAPC) through a C-terminal PDZ-binding motif and promotes intracellular bacterial proliferation and systemic virulence^16^. Analogous to its operonic partner PipB2, which is an established dual-system substrate, our profiling confirms SseM as a translocated effector with dual SPI-1/SPI-2 dependence, with translocation increasing strongly during late-stage infection, consistent with its reported role in modulating systemic growth and virulence in macrophages^16^. Notably, the relative contributions of SPI-1 and SPI-2 to cumulative effector delivery varied widely among dually translocated effectors. For example, SseM exhibited an estimated SPI-1 contribution of ∼44% of total WT translocation, whereas for SteC and SseJ the SPI-1–dependent fraction accounted for only ∼5% of total signal (**Fig. 6, Supplementary Fig. S4** and **Supplementary Table S5**). Nevertheless, for effectors with high overall translocation levels, such as SseJ, even small relative SPI-2 contributions correspond to substantial absolute amounts of effector delivery, supporting potential functional relevance despite low proportional values (**Fig. 8**).

Canonical invasion-associated effectors such as SopB, SopE, and SipA were delivered at markedly high levels during mid-stage infection, coinciding with the second wave of SPI-1 activity and rapid intracellular bacterial replication, whereas effectors involved in host signaling modulation, including SseK glycosyltransferases and the deubiquitinase SseL, were translocated at lower, yet statistically significant levels (**Fig. 8** and **Supplementary Table S5**). Although luminescence gradually decreases over time, even in the presence of the stabilized substrate vivazine, relative signal intensities across effectors and strain backgrounds provide a quantitative proxy for comparative translocation efficiency at each time point.

## DISCUSSION

The pathogenicity of *Salmonella enterica* serovar Typhimurium is fundamentally shaped by its ability to orchestrate the delivery of a diverse suite of effector proteins into host cells via two distinct T3SSs. These macromolecular injectisomes, encoded within the *Salmonella* Pathogenicity Islands, were historically conceptualized as temporally segregated systems, with T3SS-1 driving epithelial invasion and early host modeling, and T3SS-2 facilitating intracellular survival and replication within the SCV. However, accumulating experimental evidence has challenged this binary model, revealing substantial spatiotemporal overlap and functional interdependence between these two T3SSs. Here, by tracking effector translocation in real-time during the post-invasion intracellular stage of epithelial infection (2-24 hpi) using NanoBiT complementation, we generated a quantitative, system-wide map of when and through which T3SS each of the 39 annotated SL1344 effectors is delivered. A central outcome of this analysis is the extent of functional overlap between SPI-1 and SPI-2 secretion. Nearly half of the translocated repertoire showed statistically supported contributions from both systems under our experimental conditions, expanding and quantifying earlier reports of dual-system substrates. Notably, several effectors commonly considered SPI-2–biased—including SopD2, SseL, and the recently identified effector SseM—displayed measurable T3SS-1–dependent translocation during the early-to-mid intracellular phase of infection. These findings indicate that dual delivery is more prevalent than typically appreciated when assessed comprehensively and longitudinally, and that SPI-1 can contribute to activities traditionally associated with SPI-2 earlier in intracellular infection, suggesting broader functional redundancy or synergism between the two secretion systems than previously recognized.

In epithelial cells, where cytosolic hyper-replicating and vacuolar bacterial populations can develop in parallel—a phenotype not restricted to in vitro models but also observed in vivo^31^—this coexistence may help ensure effective effector deployment across distinct intracellular niches and stages of infection. A clear temporal stratification of effector delivery was observed, encompassing an early intracellular post-invasion phase dominated by SPI-1–associated effectors (2-6 hpi), a secondary T3SS-1 wave coinciding with cytosolic hyper-replication (8-10 hpi), and a later phase characterized by T3SS-2–dependent effectors that support SCV maturation and maintenance, indicating that distinct effector cohorts are deployed in coordination with transitions between intracellular lifestyles. The second T3SS-1 wave indicates that SPI-1 activity is not restricted to epithelial entry but can re-emerge after invasion to support intracellular proliferation. Consistent with this, T3SS-1—but not T3SS-2—has been shown to be required for delivery of the effectors SopB and SipA by cytosolic hyper-replicating *Salmonella*, directly linking SPI-1 activity to the cytosolic intracellular lifestyle^13^. This model is further supported by prior reports of late-phase SPI-1 effector activity, including SopB and PipB2, during intracellular infection^13 31, 46^. Notably, PipB2 has been shown to reach maximal translocation levels at approximately 16 hpi in epithelial cells^46^, closely matching our observation that overall SPI-1 effector translocation peaks around ∼17 hpi within the 2–24 hpi window.

Interestingly, in SPI-2-deficient (Δ*ssaV*) strains, dual-system effectors exhibited earlier T_max_ values than SPI-1-exclusive effectors, suggesting preferential deployment during the early intracellular phase of infection. This shift may reflect translocation from bacteria residing within nascent or destabilized SCVs, where residual SPI-1 activity persists or from cytosolic bacteria during the second wave of SPI-1 expression.

Earlier work has reported apparent hierarchies among invasion effectors, inferred either from differential secretion kinetics or from faster intracellular turnover of specific effectors^51, 52^, indicating that temporal control has often been deduced from effector function or persistence rather than from direct measurements of delivery. In contrast, our population-based real-time analysis reveals hierarchy at the level of effector translocation itself, showing that overall SPI-1-exclusive effector delivery peaks during the hyper-replication phase and that cumulative translocation across the 2–24 hpi window is strongly skewed toward a small subset of effectors. One potential mechanistic layer contributing to this hierarchy may lie in effector-specific chaperone interactions, as type III secretion chaperones are required for efficient engagement of their cognate effectors with the secretion apparatus and subsequent translocation^53, 54^. Notably, InvB-associated effectors—including SipA, SopE, SopE2, and SopA^55, 56^—cluster tightly in their peak translocation timing (T_max_ ≈ 9 hpi), consistent with a model in which chaperone–effector complexes contribute to prioritized or coordinated engagement with the T3SS during intracellular infection^53, 57^.

Although our experimental system does not distinguish vacuolar from cytosolic bacteria, future studies combining translocation reporters with intracellular localization markers, specific mutants, or differential antibiotic protection assays should enable more precise resolution of niche-specific effector deployment dynamics. Independent of spatial resolution, additional normalization strategies—such as correction for bacterial load or reference strains—may further improve quantitative comparisons of effector deployment across conditions and genetic backgrounds.

While the HiBiT-tagging approach enabled physiologically relevant expression and highly sensitive endogenous detection, tagging-associated artifacts were not negligible. While HiBiT fusions have previously been shown to permit secretion of effectors or T3SS regulators in cases where NanoLuc or GFP tagging interfered with their export, highlighting the advantage of minimal tags^11^, compatibility must nevertheless be assessed empirically for each protein. In our dataset, SifA showed no detectable translocation signal, most likely due to disruption of its C-terminal prenylation motif (CaaX), which is essential for membrane association and biological activity^58^. In addition, consistent with earlier reports on tagged translocon proteins^36, 37^, HiBiT-tagging of the translocon components SipB and SipC resulted in partial impairment of secretion or function, highlighting the sensitivity of translocon assembly to tagging. Together, these observations underscore the necessity of strategic tag placement—typically C-terminal for T3SS effectors to avoid disrupting N-terminal secretion signals—and of case-by-case validation when interpreting data from tagged proteins.

Compared with conventional endpoint or blot-based assays, which are time-consuming, less sensitive, and at best semi-quantitative, the split NanoLuc system, combined with the vivazine substrate, enabled dynamic and quantitative monitoring of effector translocation over extended infection periods. Translocation profiles obtained using furimazine at later time points were consistent with those measured using vivazine, supporting the robustness of the observed trends. Vivazine provides an effective signal half-life of approximately 24 hours, permitting continuous measurements across the 2–24 hpi window; however, gradual substrate depletion and declining luminescence over time cannot be fully excluded as contributors to subtle kinetic distortions. Importantly, late-peaking effectors—such as SseJ, which showed maximal relative translocation between ∼13.5 and 19.5 hpi—achieved substantial cumulative translocation, accounting for approximately 11% of total detected effector signal. By contrast, although endurazine is reported to provide signal stability for up to ∼72 hours, its markedly lower brightness limited sensitivity and precluded systematic detection across the effector repertoire under our experimental conditions (data not shown).

Across the 2-24 hpi window, SPI-2–exclusive translocation events accounted for only ∼1% of the total translocation signal, underscoring the quantitative dominance of SPI-1–associated delivery in this epithelial infection model. Importantly, our approach enabled, for the first time, relative quantification of (cumulative) effector abundance across the translocated effector repertoire, revealing a highly skewed distribution of effector contribution: SopE (40%), SptP (19%), and SseJ (11%) together accounted for approximately 70% of the total translocation signal. This observation highlights how system-wide quantitative approaches can uncover functional weighting within the effector repertoire and suggests that the skewed distribution may reflect differences in stoichiometric and mechanistic requirements among effectors, with some activities producing strong downstream effects at low abundance, whereas effectors involved in structural remodeling or sustained signaling interference may require higher cumulative delivery. In this context, SopE and SptP have long been recognized as central SPI-1 substrates due to their prominent roles in actin remodeling and invasion dynamics^2^. Their strong early engagement has made them frequent focal points of mechanistic studies, consistent with our observation that they dominate the cumulative translocation signal when viewed across the full effector repertoire.

HiBiT blotting, while useful for endpoint validation, lacks the temporal resolution and sensitivity of the real-time translocation assay, particularly for low-abundance or late-translocated effectors. Related split-luciferase approaches have been applied to study T3SS activity (including translocon assembly) or T4SS-mediated translocation in *Shigella, Pseudomonas, Legionella*, and *Yersinia*, in addition to *Salmonella*^*11, 12, 59, 60*^, but these studies have generally focused on only a limited number of proteins or effectors.

Whereas most translocation studies in epithelial models emphasize the invasion phase, later stages have typically been examined through effector localization or single-cell imaging rather than quantitative translocation assays. Consequently, effector delivery during prolonged intracellular infection has not previously been systematically quantified across the effector repertoire. Our approach instead captures a previously undercharacterized mid-to-late intracellular window, enabling population-level measurement of effector translocation kinetics associated with intracellular niche adaptation.

In this context, our dataset—encompassing 39 effectors across multiple genetic backgrounds— provides an unprecedented systems-level view of T3SS-dependent effector delivery during epithelial infection.

Notably, several effectors (CigR, SpvB, SpvC, SpvD, SseI, SteB) showed no detectable translocation in our HeLa model. The absence of translocation signal for CigR—despite chromosomal tagging—along with its prior characterization as an inner membrane protein lacking a secretion signal^61^, argues against its continued classification as a T3SS effector in SL1344, refining the functional effector set to 38. For SseI and Spv family members, the lack of signal likely reflects HeLa-specific regulation, as these effectors play roles predominantly in systemic infection and macrophage survival^42^. These observations illustrate the context dependency of effector deployment and highlight the value of host-specific models^44^.

Although effector translocation profiles were generally reproducible across experiments, SPI-2 induction is known to be more heterogeneous^62^, and this variability may contribute to an increased dispersion of late-stage translocation kinetics. Except for the prophage-associated effector SseK3, which showed comparatively earlier translocation (T_max_ ∼15 hpi), T_max_ values for SPI-2 effectors clustered more closely together than those of SPI-1 effectors, suggesting a more synchronized deployment once SPI-2 is activated. Together, the resurgence of SPI-1 activity during the second wave coinciding with cytosolic hyper-replication and the delayed yet coordinated delivery of SPI-2 effectors (e.g., SseF, SseG, and SseK1–3) support a model in which intracellular persistence is maintained through temporally tuned engagement of both secretion systems, consistent with coexisting vacuolar and cytosolic subpopulations rather than a strict sequential switch from SPI-1 to SPI-2.

Temporal regulation of secretion may occur at multiple levels—transcriptional control, chaperone availability, or signal recognition—and is likely subject to host-imposed cues. Together, these findings indicate that effector deployment is tightly coupled to dynamic transitions between intracellular lifestyles, rather than being governed by a single unidirectional secretion program. This study reframes the effector landscape of *Salmonella* infection as a temporally dynamic continuum characterized by overlapping contributions from both T3SSs. By mapping effector delivery kinetics at high temporal resolution, we provide a resource for identifying stages of infection that may be particularly susceptible to intervention, including the mid-phase cytosolic proliferative burst that is often overlooked. The split-NanoLuc platform is readily adaptable to broader applications—including other serovars, host cell types, or co-infection models—eanbling systematic comparisons of secretion dynamics across infection contexts. Such extensions across serovars or host tissues may reveal differential weighting or timing of effector deployment, potentially uncovering virulence strategies tuned to host range or disease outcome^15^. Notably, apparent discrepancies in the contribution of SPI-2 effectors across Salmonella infection models have been attributed to short infection windows and the limited sensitivity of bulk c.f.u. measurements^15^, highlighting the need for extended, time-resolved analyses that simultaneously capture effector translocation and intracellular bacterial replication—an advance enabled by the combined translocation and GFP-based replication readouts assessed here.

In conclusion, our findings challenge a strictly segregated view of T3SS function and instead support a model in which effector translocation is continuous, flexible, and dynamically regulated throughout intracellular infection. This refined understanding of secretion dynamics, combined with scalable and sensitive detection platforms, provides a framework for dissecting how bacterial pathogens coordinate complex virulence programs in real-time. Ultimately, integrating high-resolution kinetic maps of effector delivery with biochemical, structural, and single-cell approaches will be essential for elucidating how *Salmonella* seamlessly navigates the complex intracellular landscape of the host to establish a permissive niche for survival and replication.

## METHODS

### Bacterial strains and culture conditions

The *Salmonella enterica* serovar Typhimurium (*S*. Typhimurium) wild-type strain SL1344^63^ (Genotype: *hisG46*, Phenotype: His(-); biotype 26i) was obtained from the *Salmonella* Genetic Stock Center (SGSC, Calgary, Canada; cat. no. 438^63^). A GFP-expressing SL1344 strain (JVS-3858), carrying P_tet_ ::*gfp* integrated into the chromosomal *putPA* locus^27^, was kindly provided by Prof. Jörg Vogel and Prof. Alexander Westermann (**Supplementary Table S3**).

Bacterial cultures were grown either in LB Miller broth (10 g/L Bacto tryptone, 5 g/L Bacto yeast extract, 10 g/L NaCl) or under SPI-2–inducing conditions in LPM minimal medium (5 mM KCl, 7.5 mM (NH_4_)_2_SO_4_, 0.5 mM K_2_SO_4_, 38 mM glycerol (0.3% v/v), 0.1% casamino acids, 8 μM MgCl_2_, 337 μM PO_4_ ^3-^, and 80 mM MES, pH 5.8)^19^.

For routine culturing, single colonies were picked from LB agar plates supplemented with the appropriate antibiotics, inoculated into 2 mL LB broth in round-bottom culture tubes (Corning, cat no. 352059), and grown for ∼18 h at 37°C with agitation (180 rpm). For LB experiments, O/N cultures (OD_600_ ∼ 2.0–2.5) were diluted to OD_600_ 0.02 in antibiotic-free LB in ventilated T25 flasks (8 mL per culture) or in 24-well plates (Greiner, cat. no. 662102) sealed with gas-permeable porous tape (Micropore^TM^ surgical tape, 1-inch width; 3 M, product no. 1530). Cultures were incubated at 37°C and 180 rpm until the early stationary phase (OD_600_ ∼ 2.0) or grown O/N, as indicated. For SPI-2 induction, O/N LB cultures were washed twice in LPM medium and reinoculated at OD_600_ 0.02 into fresh LPM medium. Cultures were grown at 37°C with shaking (180 rpm) for ∼6 h until reaching OD_600_ ∼ 0.6. All strains used in this study are listed in **Supplementary Table S1**.

### Plasmids

Plasmids pKD46 (#14175) and pKD4 (#7632) were obtained from the Coli Genetic Stock Center (CGSC, Yale University, USA). Plasmid pKD3 (#45604) and the recombineering helper plasmid pORTMAGE-2^64^ (#72677) were obtained from Addgene, and plasmid pDL17^65^ was kindly provided by Dr. Dmitrii Bubnov and Prof. Sergey Sineoky. Plasmids pKD3 and pKD4 are π-dependent suicide vectors encoding chloramphenicol (Cam^R^) and kanamycin (Kan^R^) resistance cassettes, respectively^17^. All three helper plasmids (pKD46^17^, pORTMAGE-2^64^, and pDL17^65^) encode the bacteriophage λ Red recombineering genes (*γ, β*, and *exo*), which catalyze homologous recombination using linear double-stranded DNA in bacteria^17^. Plasmid pKD46 expresses the λ Red system under the arabinose-inducible P_BAD_ promoter and carries a temperature-sensitive *repA* origin^17^. The pORTMAGE-2 plasmid additionally encodes a dominant-negative *mutL*(E32K) allele to increase recombineering efficiency; expression of its recombineering components is controlled by the temperature-sensitive cI857 repressor^64^. Plasmid pDL17 likewise has λ Red functions driven by the P_*rhaB*_ promoter^65^.

### λ Red recombineering for mutant and HiBiT-tagged strain construction

Mutant and HiBiT-tagged *S*. Typhimurium SL1344 strains were generated by λ Red recombineering, as previously described^17^ (**Supplementary Tables S2-3**). Cam^R^ and Kan^R^ resistance cassettes were PCR-amplified from plasmids pKD3 and pKD4, respectively, using primers listed in **Supplementary Table S4**^17^. Oligonucleotide primers were synthesized by Integrated DNA Technologies (IDT, Leuven, Belgium). All recombineering PCR primers contained a 50 bp 5’ homology extension targeting the genomic region of interest, flanking either the full coding sequence or, in the case of the *ssaV* deletion (SL1344 Δ*ssaV::cat*), a defined internal segment as previously described^66^. For C-terminal tagging, homology arms flanked the 3⍰ coding region. For HiBiT-tagging, the forward primer additionally encoded the 33 bp *HiBiT* sequence immediately followed by the native stop codon of the effector gene of interest. Single-stranded DNA oligonucleotides for oligo-mediated allelic replacement (OMAR) were designed with the Mage Oligo Design Tool^67^ (MODEST; ‘translational_knockout’ operation) and were multiplexed with PCR-derived, double-stranded linear editing substrates for recombineering, as previously described^68, 69^. PCR-derived linear editing substrates were (co-)electroporated (together with the corresponding OMAR single-stranded DNA oligonucleotides) into SL1344 strains harbouring one of the recombineering helper plasmids pKD46^17^, pDL17^65^, or pORTMAGE-2^64^. Electrocompetent cells were prepared from cultures grown to OD_600_ ∼ 0.6. Expression of λ Red recombination functions was induced according to the respective helper plasmid system: L-arabinose induction (0.2% w/v) for pKD46; L-rhamnose induction (0.2% w/v L-rhamnose) for pDL17; or heat-induction at 42°C for pORTMAGE-2, as described^17, 64, 65^. Approximately 2.4 × 10^9^ induced electrocompetent cells were electroporated with the linear PCR-derived editing substrates encoding the Kan^R^ or Cam^R^ cassette. Immediately following electroporation, cells were recovered in pre-warmed Super Optimal broth with Catabolite repression (SOC; 2% tryptone, 0.5% yeast extract, 10 mM NaCl, 2.5 mM KCl, 5 mM MgCl_2_, 10 mM MgSO_4,_ and 20 mM glucose) with shaking (180 rpm) either at 37°C for 2-3 h (pKD46 and pDL17) or at 28°C for 3-4 h (pORTMAGE-2) before plating on LB agar supplemented with kanamycin (50 µg/mL; Duchefa Biochemie, cat. no. K0126) or chloramphenicol (25 µg/mL; Sigma-Aldrich, cat. no. C0378). Plates were incubated O/N at 37°C (pKD46- and pDL17-derived recombinants) or 28°C (pORTMAGE-2-derived recombinants). Following re-streaking, correct allelic replacement or C-terminal HiBiT insertion was verified by colony PCR using locus-specific primers (**Supplementary Table S4**), and PCR products were confirmed by Sanger sequencing (Eurofins Genomics).

### P22 Phage transductions

P22 HT-mediated transductions were performed using a high-titer P22 lysate (>10^10^ PFU/mL) propagated on *Salmonella enterica* serovar Typhimurium SGSC1412 (LT2 prototroph), obtained from the *Salmonella* Genetic Stock Center (Calgary, Canada; cat. no. 496), as previously described^70^. Transductions were used to introduce defined genetic backgrounds into *S*. Typhimurium SL1344 strains carrying endogenously HiBiT-tagged effectors (Kan^R^). High-titer P22 HT lysates were generated from SL1344 Δ*invA::cat* (PVD0316), SL1344 Δ*ssaV::cat* (PVD0317), and SL1344 P_tet_::*gfp cat* (PVD0002) donor strains following standard procedures^71^. Prepared lysates were used to infect the endogenously HiBiT-tagged effector strains, and transductants were selected on LB agar supplemented with kanamycin (50⍰µg/mL) and chloramphenicol (25⍰µg/mL), as well as 2⍰mM EGTA and 5⍰mM sodium citrate to suppress phage propagation. After restreaking to eliminate residual phage, successful transductants were confirmed by colony PCR using locus-specific primers (**Supplementary Table S4**). The resulting strain panel, comprising HiBiT-tagged effectors in Δ*invA*, Δ*ssaV*, or P_tet_::*gfp* backgrounds, is detailed in **Supplementary Table S2**.

### Mammalian cell culture

HeLa cells (human cervical adenocarcinoma, American Type Culture Collection, Manassas, VA, USA; ATCC*®* CCL-2™) and a stable HeLa cell line expressing LgBiT (HeLa-LgBiT)^11^, kindly provided by Prof. Samuel Wagner, were maintained in GlutaMAX^TM^-supplemented Dulbecco’s Modified Eagle Medium (DMEM; Invitrogen, cat. no. 31966047) containing 10% foetal bovine serum (FBS; Invitrogen, cat. no. 10270106) and antibiotics (50 units/mL penicillin and 50 µg/mL streptomycin; Gibco, cat. no. 15070063). For infection experiments, cells were switched to antibiotic-free medium supplemented with heat-inactivated FBS (30 min at 56°C). For live-cell real-time imaging in 96-well plates (see below methods), culture medium was filter-sterilized using 0.22 µm filters. All cells were grown at 37°C in a humidified 5% CO_2_ atmosphere and routinely passaged every 3–4 days.

### Bacterial total protein extraction

Bacterial cultures, grown O/N or harvested at defined growth phases, were collected by centrifugation at 6,000 × g for 10 min at 4°C, and supernatants were discarded. Bacterial pellets were washed once with ice-cold Dulbecco’s phosphate-buffered saline (DPBS; Gibco, cat. no. 14190144), transferred to 1.5 mL microcentrifuge tubes, and centrifuged again at 6,000 × g for 5 min at 4°C. Supernatants were removed, and pellets were stored at -80°C until further processing. Frozen bacterial pellets were resuspended in urea lysis buffer (9 M urea, 50 mM ammonium bicarbonate (NH_4_HCO_3_), pH 7.9) to obtain a final protein concentration of approximately 2–3 mg/mL^72^. Cell lysis was performed by three cycles of freezing in liquid nitrogen and thawing at room temperature, followed by two cycles of microprobe sonication (LABSONIC® M, type BBI 853 5027) using a 1-mm probe at 50% duty cycle, 0.5-s pulses for 30 s, and an amplitude of 40. Insoluble debris was removed by centrifugation at 16,000 × g for 10 min at 4°C, and clarified lysates were collected. Protein concentrations were determined using the DC^TM^ Protein Assay Kit (Bio-Rad, cat. no. 5000006) according to the manufacturer’s instructions.

### SDS–PAGE, HiBiT blotting, and immunoblotting

Protein samples for one-dimensional SDS-PAGE were prepared by the addition of XT sample loading buffer (4×; Bio-Rad, cat. no. 1610791) and XT reducing agent (20×; Bio-Rad, cat. no. 1610792) according to the manufacturer’s instructions. Precision Plus Protein™ All Blue Prestained Protein Standards (Bio-Rad, cat. no. 1610373) were used as molecular weight markers. Proteins were resolved on 4–12% gradient or 12% XT precast Criterion gels (Bio-Rad) using XT-MOPS running buffer (Bio-Rad, cat. no. 1610788) at 150–200 V and transferred to 0.45-µm nitrocellulose membranes for 30 min at 100 V using Tris–borate transfer buffer (50 mM Tris (base) and 50 mM boric acid). Membranes were rinsed for 10 min in Tris-buffered saline containing 0.1% Tween-20 (TBS-T; 50 mM Tris-HCl, 150 mM NaCl, 0.1% (v/v) Tween-20, pH 7.5) and subsequently processed for HiBiT detection or immunoblotting as described below.

For HiBiT blotting, membranes were incubated with recombinant LgBiT protein (Promega, cat. no. N401C; 1:200 dilution) or in-house–produced recombinant 6×His-LgBiT protein^73^ (final concentration 25 nM) in TBS-T for 1 h at room temperature or O/N at 4°C. Chemiluminescent detection was performed by incubation with Nano-Glo® Luciferase Assay Substrate (Promega, cat. no. N113A; 1:2000 dilution) or furimazine substrate (TargetMol, cat. no. T15359; final concentration 5 nM) in TBS-T for 5 min under gentle agitation.

For immunoblotting, membranes were blocked for 30 min at room temperature in a 1:1 mixture of TBS-T and Odyssey blocking buffer (LI-COR, cat. no. 927-40003) and incubated overnight at 4°C with primary antibodies diluted in the same buffer. Membranes were washed three times for 5 min in TBS-T and incubated with fluorescent secondary antibodies, diluted in a 1:1 TBS-T-Odyssey blocking buffer mixture, for 30 min at room temperature, followed by three washes in TBS-T and a final wash in TBS.

The following primary antibodies and affinity reagents were used: mouse anti-GAPDH (Abcam, ab9484; 1:10,000), rabbit anti-GFP (Invitrogen, A11122; 1:1,000), and streptavidin–Alexa Fluor^TM^ 680 conjugate (Invitrogen, S32358; 1:5,000). Secondary antibodies were IRDye 800CW goat anti-mouse IgG (LI-COR, cat. no. 926-32210; 1:10,000) and IRDye 800CW goat anti-rabbit IgG (LI-COR, cat. no. 926-32211; 1:10,000). Endogenously biotinylated proteins detected using the streptavidin–Alexa Fluor^TM^ 680 conjugate served as loading controls.

Fluorescent (600 nm, 700 nm, and 800 nm) and chemiluminescent signals were acquired using the Odyssey Fc imaging system (LI-COR Biosciences, model no. 2800). Signal quantification was performed using Empiria Studio software (LI-COR), with background subtraction based on user-defined regions. Quantified values were exported as background-corrected signal intensities.

### Protein secretion analysis

For analysis of secreted proteins, *S*. Typhimurium SL1344 strains were cultivated in 1.5 mL LB Miller broth in 24-well plates (Greiner, cat. no. 662102) sealed with gas-permeable porous tape (Micropore^TM^ surgical tape, 1-inch width; 3 M, product no. 1530). Cultures were inoculated from O/N cultures to an initial OD_600_ of 0.02 and grown at 37°C with agitation (180 rpm) either until the early stationary phase (OD_600_ ∼ 2.0) or O/N, as indicated. Culture volumes of 1.2 mL were transferred to 1.5 mL microcentrifuge tubes, and bacterial cells were pelleted by centrifugation at 6,000 × g for 5 min at 4°C. A culture supernatant volume of 1 ml was carefully transferred to a new tube, while residual medium was removed and discarded. Bacterial pellets were retained on ice or stored at -80°C for subsequent total lysate preparation. Supernatants were passed through 0.22-µm pore-size filters to remove residual bacteria and immediately placed on ice. Proteins were precipitated by the addition of trichloroacetic acid (TCA) to a final concentration of 10% (v/v), followed by gentle mixing and incubation at 4°C overnight. Precipitated proteins were collected by centrifugation at 16,000 × g for 30 min at 4°C. The supernatant was discarded, and protein pellets were washed once with 300 µL ice-cold acetone to remove residual acid, followed by centrifugation at 16,000 × g for 30 min at 4°C. Protein pellets were air-dried for 10–15 min and resuspended by thorough vortexing in 80 µL of 50 mM Tris-HCl (pH 8.0) supplemented with SDS sample loading buffer (Bio-Rad XT). Samples were heated at 70°C for 10 min prior to analysis. Corresponding bacterial pellets were lysed in urea lysis buffer (9 M urea, 50 mM ammonium bicarbonate (NH_4_HCO_3_), pH 7.9) and processed in parallel as described above. Secretome and total lysate samples (30 µg) were resolved by SDS–PAGE and analysed by HiBiT blotting as described above. Where indicated, fivefold higher equivalents of secretome samples were analysed to enhance detection sensitivity. All secretome analyses shown are representative of at least two independent biological replicates.

### Real-time multimodal live-cell infection setup

HeLa LgBiT cells were seeded at 1.25–1.5 × 10^4^ cells per well into black, flat-bottom 96-well plates (Tecan, cat. no. 30122306) in 100 μL DMEM containing 10% heat-inactivated FBS, and incubated 24 h prior to infection at 37°C in a humidified 5% CO_2_ incubator. For infection, *S*. Typhimurium SL1344 cultures were grown O/N in LB, diluted to an OD_600_ of 0.02, and incubated at 37°C with shaking until reaching late logarithmic/early stationary phase (OD_600_ ∼ 3.0; ∼2 × 10^9^ CFU/mL; ∼4 h growth). This physiological stage is associated with maximal SPI-1 induction and epithelial invasion competence^74^. Cells were harvested by centrifugation (6,000 × g, 5 min, 20°C), washed once in Hank’s balanced salt solution (HBSS; +Ca^2+^/Mg^2+^; Gibco, cat. no. 14025100), re-centrifuged, and resuspended in HBSS to the required density. Fifty microliters of bacterial suspension were added per well to obtain the indicated multiplicity of infection (MOI 10–100), and infection proceeded for 30 min at 37°C. Monolayers were then aspirated and washed once with pre-warmed HBSS (200 μL/well). To eliminate extracellular bacteria, monolayers were overlaid with 150 μL DMEM containing 10% heat-inactivated FBS and 100 μg/mL gentamicin (Sigma, cat. no. G1264-1G) for 1 h at 37°C. Subsequently, medium was replaced with 150 μL DMEM containing 10% heat-inactivated FBS, 10 μg/mL gentamicin, and Nano-Glo® Vivazine™ Live Cell Substrate (Promega, cat. no. N2580; final dilution 1/150), pre-equilibrated for ∼1 h at 37°C to enable substrate activation.

### Real-time live-cell luminescence, fluorescence, and imaging acquisition

Real-time acquisition of the live-cell infection plates was initiated 2 hpi and continued until 26 hpi using a Spark Cyto 400 multimodal plate reader (Tecan) operated through SparkControl™ (v3-4; Tecan, Mechelen, Belgium) software. Prior to imaging, plates were sealed with sterile, gas-permeable, optical adhesive sealing film and placed inside the instrument’s small humidity cassette. During acquisition, plates were maintained at 37°C in a humidified atmosphere containing 5% CO_2_ via the integrated environmental control system.

Bright-field images were captured every 30 min using a 10x objective, acquiring a single centred field per well. Automated confluence and cell-count analyses were performed on these images using the Image Analyzer™ module (Tecan), with post-acquisition sensitivity adjustments applied when required. Whole-well luminescence was recorded every 15–30 min using automatic attenuation, a 0-ms settle time, and a 1,000-ms integration period. Where applicable, eGFP fluorescence signals were recorded at matching intervals to monitor bacterial load dynamics using bottom-reading optics (excitation: 485 nm ± 20 nm; emission: 535 nm ± 20 nm). Fluorescence analysis was performed using segmentation-based settings (sensitivity 70%; object length/width 8–30 µm). Each strain and infection condition was assayed in 3–4 technical replicates per experiment.

### Effector translocation quantification and statistical analysis

Luminescence time-course data were quantified by calculating the area under the curve (AUC); expressed as relative light units x hours (RLU.h) following background subtraction. AUC values were calculated by trapezoidal numerical integration (i.e., summing the areas of trapezoids defined by consecutive time points and their corresponding luminescence values) over defined infection intervals: overall (2–24 hpi), mid-stage infection (2–10 hpi; corresponding to the second wave of T3SS1 activity), and late-stage infection (11–24 hpi). AUC values were computed for each biological replicate in WT (eGFP+), Δ*invA*, and Δ*ssaV* strain backgrounds. For SipA, the corresponding HiBiT-tagged Δ*invA* strain was not available; therefore, Δ*invA* background values from the *cigR::cigR-HiBiT* Δ*invA* strain measured on the same plate were used for AUC calculations.

Effectors with WT mean overall AUC < 200 were considered not detectably translocated. For SifA, which showed elevated baseline luminescence without a clear infection-associated increase, a more stringent overall AUC threshold of 300 was applied. For phase-specific analyses, a WT mean AUC threshold of 150 was applied to ensure measurable signal during the corresponding interval. Statistical comparisons were performed separately for mid-stage and late-stage infection phases using one-way analysis of variance (ANOVA) followed by pairwise Welch’s t-tests. P-values were adjusted for multiple testing using the Benjamini–Hochberg false discovery rate (FDR) procedure independently for each phase, with FDR ≤ 0.01 considered statistically significant. Effectors were classified as SPI-1, SPI-2, or SPI-1/2 dependent based on statistically significant, phase-specific differences combined with phase-specific AUC thresholds, or as translocated without statistically supported secretion system dependence when WT signal was detected but no significant mutant-dependent differences were observed.

For effectors classified as SPI-1/2 dependent, as well as for effectors showing detectable WT translocation but lacking statistically supported secretion system dependence, the relative contribution of SPI-1 and SPI-2 to cumulative effector delivery was further quantified using overall AUC values (2–24 hpi). Mean overall AUC values were compared between WT and Δ*ssaV* strains, where Δ*ssaV* was considered to represent SPI-1–dependent delivery in the absence of SPI-2 function, and WT represented the combined contribution of SPI-1 and SPI-2. The relative SPI-1 contribution was calculated as the fraction of the WT signal retained in the Δ*ssaV* background (SPI-1 (%) = (Δ*ssaV* AUC / WT AUC) x 100). The SPI-2 contribution was defined as the complementary fraction of the WT signal (SPI-2 (%) = 100 - SPI-1 (%)).

### Host cell fractionation and isolation of intracellular bacteria

HeLa cells were seeded at 5.25 × 10^5^ cells per well in 6-well plates (Thermo Scientific, Nunclon^TM^ cat. no. 140675) one day before infection and infected at an MOI of 50 or 100, as indicated and described above. At the indicated hpi, infected cell cultures were washed with ice-cold DPBS (Gibco, cat. no. 14190094), scraped in 1 mL DPBS on ice, and transferred to 1.5 mL microcentrifuge tubes. Cells were pelleted by centrifugation at 300 × g for 10 min at 4°C and subsequently lysed in 200 µL lysis buffer containing 20 mM Tris-HCl (pH 7.6), 150 mM NaCl, and 0.1% Triton X-100. Cell lysates were centrifuged at 600 × g for 5 min at 4°C to remove nuclei and cellular debris. The resulting supernatant was transferred to a new tube and centrifuged at 4,000 × g for 20 min at 4°C to separate Triton X-100–soluble host cell fractions (containing translocated effectors) from Triton X-100–insoluble material. Triton X-100–soluble fractions were supplemented with XT sample loading buffer and XT reducing agent (according to the manufacturer’s instructions), and processed for SDS– PAGE as described above.

The Triton X-100–insoluble pellets, containing bacteria-associated material, were washed once with lysis buffer to remove residual host proteins, flash-frozen in liquid nitrogen, and stored at -80°C until further processing. Bacterial pellets were subsequently lysed in urea lysis buffer (9 M urea, 50 mM ammonium bicarbonate (NH_4_HCO_3_), pH 7.9) and processed as described above. Host cell and bacteria-associated fractions were resolved by SDS–PAGE and analysed by HiBiT blotting as described above. For SDS–PAGE analysis, 50 µg of host-cell (TX-100–soluble) protein and the corresponding bacteria-associated (TX-100–insoluble) equivalents were analysed unless stated otherwise.

### Software packages for statistics and data visualization

For basic data handling, statistical testing, calculation of summary statistics, and analysis of time-course and AUC-based data, GraphPad Prism was used (Prism 10, v10.6.0; GraphPad Software, San Diego, CA, USA; www.graphpad.com). Graphical representation of standard plots (time courses, bar graphs, scatter plots) was also performed in GraphPad Prism. For specialized data visualization, including Voronoi treemaps and matrix (heatmap-style) plots, RAWGraphs 2.0 was used (https://app.rawgraphs.io/). In addition, custom Python scripts were used for automated data parsing, phase-specific AUC calculations, peak-window determination, timing metric extraction (T_25_, T_50_, T_max_), and batch statistical comparisons across effectors and strain backgrounds.

## ACKNOWLEDGEMENTS

We thank Prof. Jörg Vogel and Prof. Alexander Westermann for providing the GFP-expressing *Salmonella* SL1344 strain JVS-3858 (P_tet_ ::*gfp*)^27^, and Prof. Samuel Wagner and Dr. Erwin Bohn for providing the stable HeLa cell line expressing LgBiT (HeLa-LgBiT)^11^, and Valdes Snauwaert for generating the *sifA* tko strain. This work was made possible through the support of the European Research Council (ERC) under the European Union’s Horizon 2020 research and innovation program (PROPHECY grant agreement no. 803972 to PVD), the Fonds Wetenschappelijk Onderzoek (FWO-Vlaanderen; project G045921N to PVD and a doctoral fellowship to L.S., grant 1119925N), and the Ghent University Concerted Research Actions (grant BOF23/GOA/001 to PVD). The funding bodies had no role in study design, data collection or analysis, data interpretation, manuscript preparation, or the decision to publish.

## AUTHOR CONTRIBUTIONS

PVD conceived the study and designed the experiments. VJ and PVD performed the experiments. PVD analyzed the data. PVD wrote the manuscript, LS contributed to Table preparations and finalizing the manuscript text, and all authors gave approval to the final version of the manuscript.

## DECLARATION OF AI USE

ChatGPT and QuillBot were used during the final preparation of this manuscript to improve language and readability. No scientific content was generated by these tools.

## SUPPLEMENTARY FIGURES

**Supplementary Figure S1.**
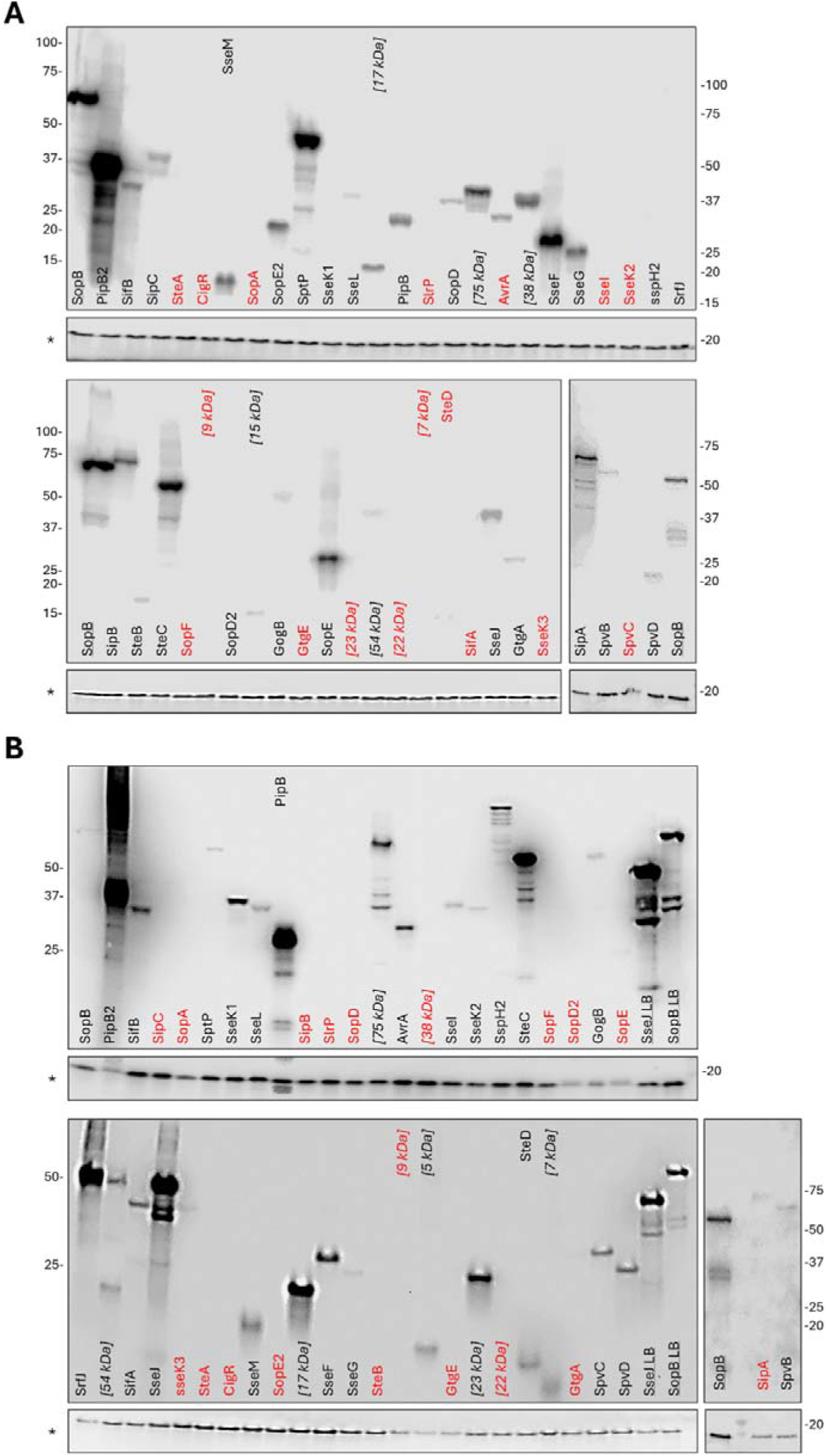
HiBiT blotting validation of endogenously HiBiT-tagged effector expression under SPI-1 and SPI-2 inducing conditions. **(A)** HiBiT blotting of total lysates (TLs) from LB overnight (O/N) cultures (SPI-1–permissive conditions). Upper panels show HiBiT blots of whole-cell lysates from Salmonella Typhimurium SL1344 strains carrying endogenous C-terminal HiBiT fusions to individual type III effector proteins. Detection was performed by HiBiT–LgBiT complementation, enabling highly sensitive luminescent detection of intact HiBiT-tagged precursors. The biotin carboxyl carrier protein (BCCP; 17 kDa), used as a loading control, was probed with a fluorescent streptavidin conjugate (marked with *). Lanes marked only with a molecular weight (MW) between square brackets represent HiBiT-tagged proteins not included among the 39 effector proteins analysed in this study and are therefore shown only for completeness of the blot layout. Red labels indicate HiBiT-tagged proteins that were not detectably expressed under the indicated condition, whereas black labels denote proteins with a detectable HiBiT signal. MW markers are shown to the left or right of each blot, as indicated. The LB O/N samples for SopB were included as internal reference controls across HiBiT blots and were used for the relative quantification of expression intensities reported in **Fig. 1**, following normalisation to the corresponding BCCP loading control. **(B)** HiBiT blotting of TLs from LPM pH 5.8 cultures (SPI-2–inducing conditions). As in panel A, HiBiT blots of TLs from HiBiT-tagged SL1344 strains grown for 6 h in mildly acidic LPM medium (pH 5.8) to OD_600_ ≈ 0.6 are shown, representing classical SPI-2 induction conditions. The biotin carboxyl carrier protein (BCCP; 17 kDa), used as a loading control, was probed with a fluorescent streptavidin conjugate (marked with *). Lanes labelled solely with MW annotations correspond to HiBiT-tagged proteins unrelated to the effector panel analysed in this study. Red and black labeling follows the same convention as in panel A. MW standards are shown on the appropriate side of each blot. SopB (LB O/N) and SseJ (LB O/N) reference samples were run on all blots to enable cross-gel normalisation (as used for the quantifications shown in **Fig. 1**), ensuring comparability of relative expression across HiBiT blots.

**Supplementary Figure S2.**
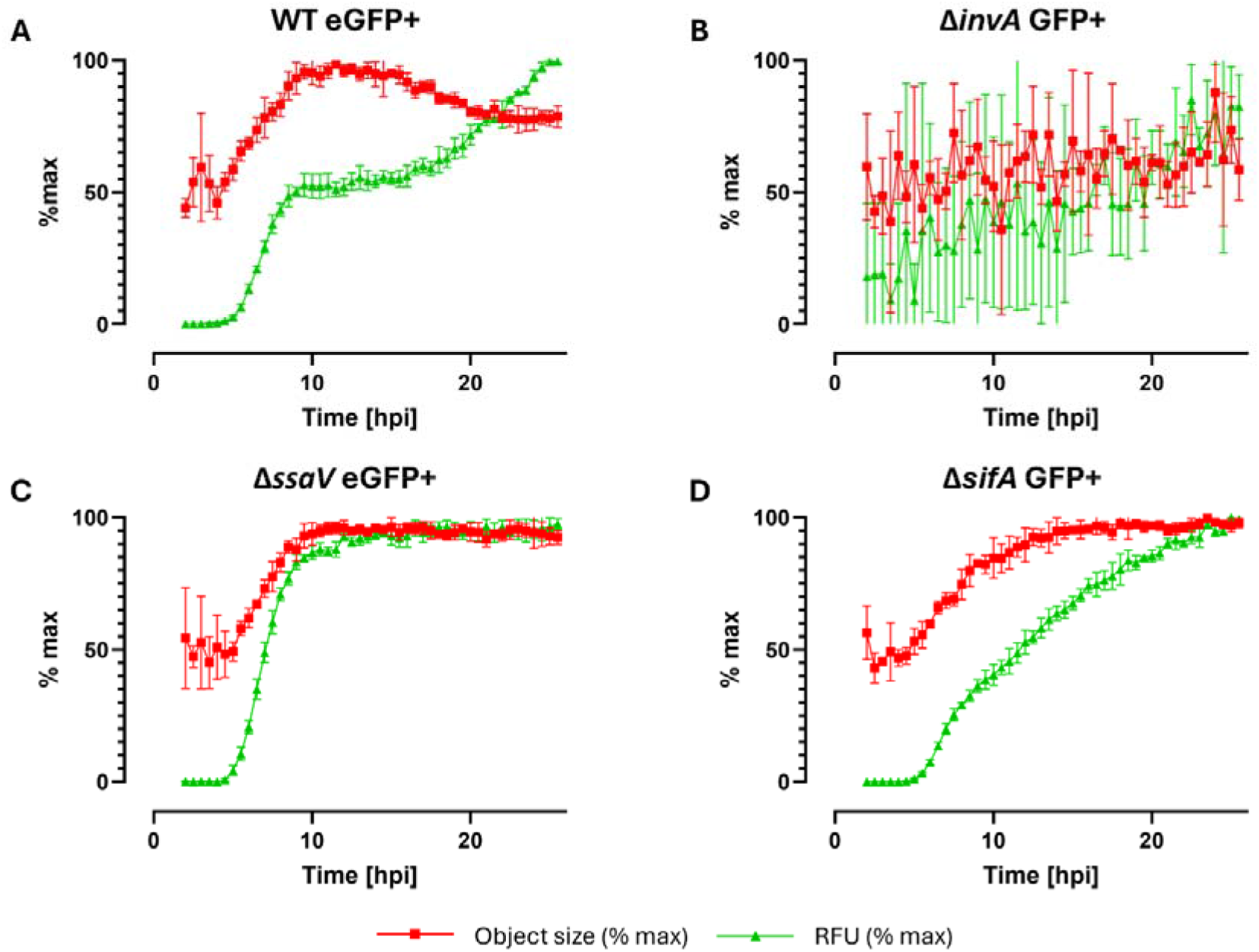
Real-time profiling of intracellular replication dynamics of wild-type and virulence-deficient *Salmonella* SL1344 strains in HeLa cells. HeLa cells were infected with constitutively eGFP-expressing S. Typhimurium SL1344 wild-type (WT) or mutant strains deficient in key virulence factors: ΔinvA (T3SS-1-deficient), ΔssaV (T3SS-2-deficient), or a translational knockout of the SPI-2 effector SifA (ΔsifA, tko). Infections were performed at a multiplicity of infection (MOI) of 50, and real-time monitoring was conducted over 24 hours using automated label-free imaging in a multimodal plate reader under environmental control. Fluorescence parameters were normalized to their respective maxima (% max). Total green fluorescence (RFU, green) and average green object size (µm^2^, red) serve as proxies for intracellular bacterial load and bacterial distribution within infected cells, respectively. Data points represent mean ± SD of technical quadruplicates.

**Supplementary Figure S3.**
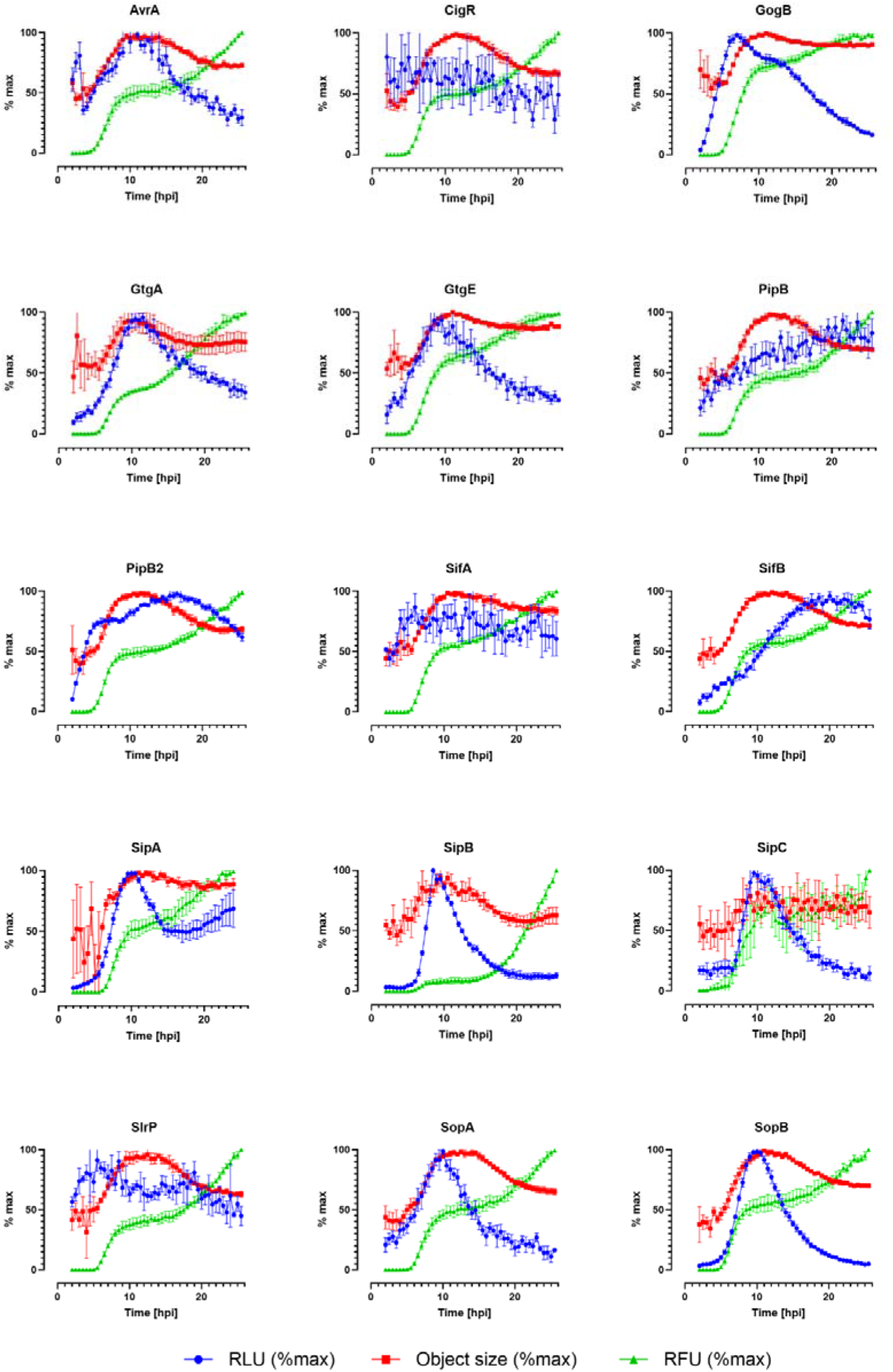
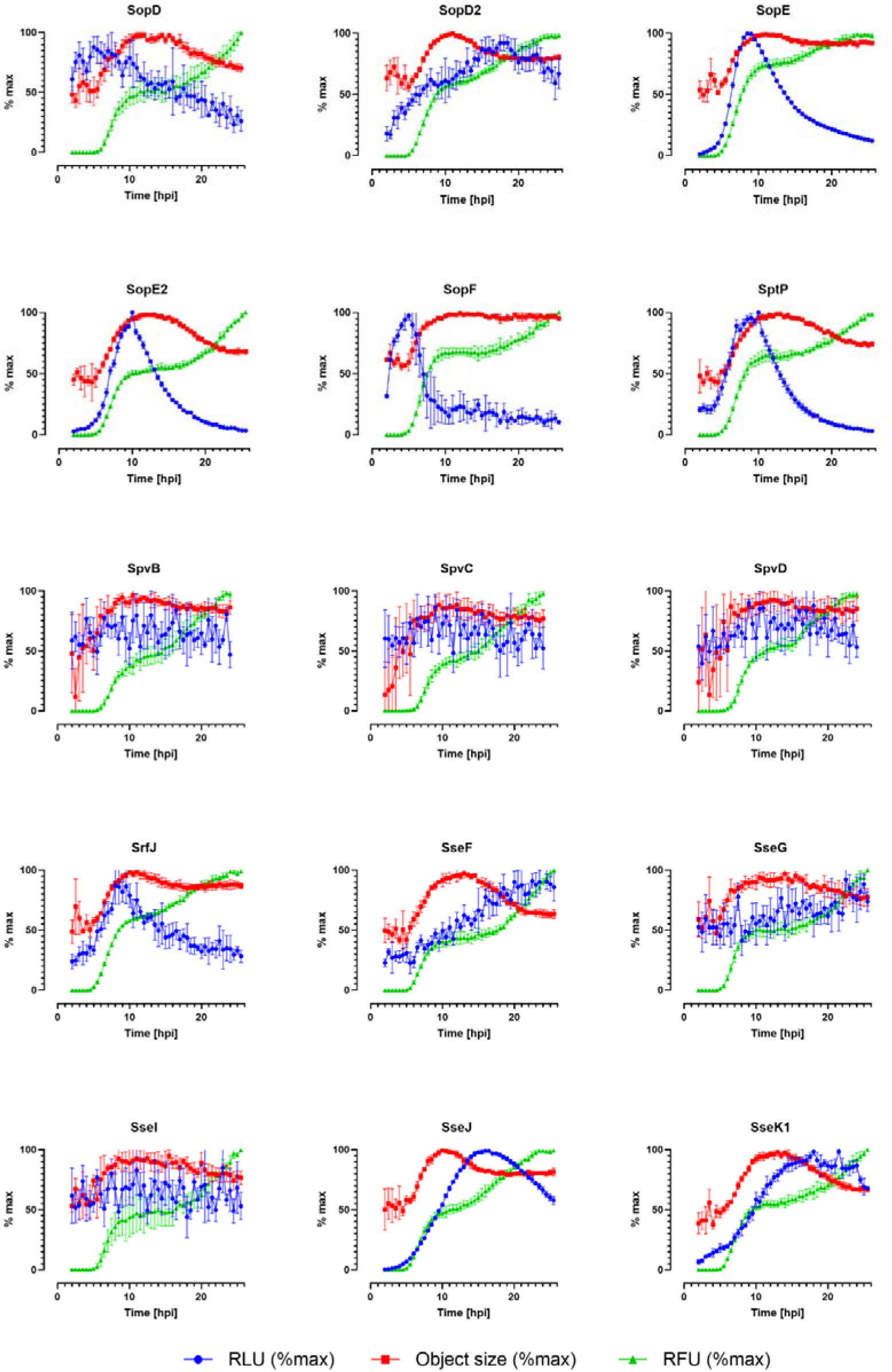
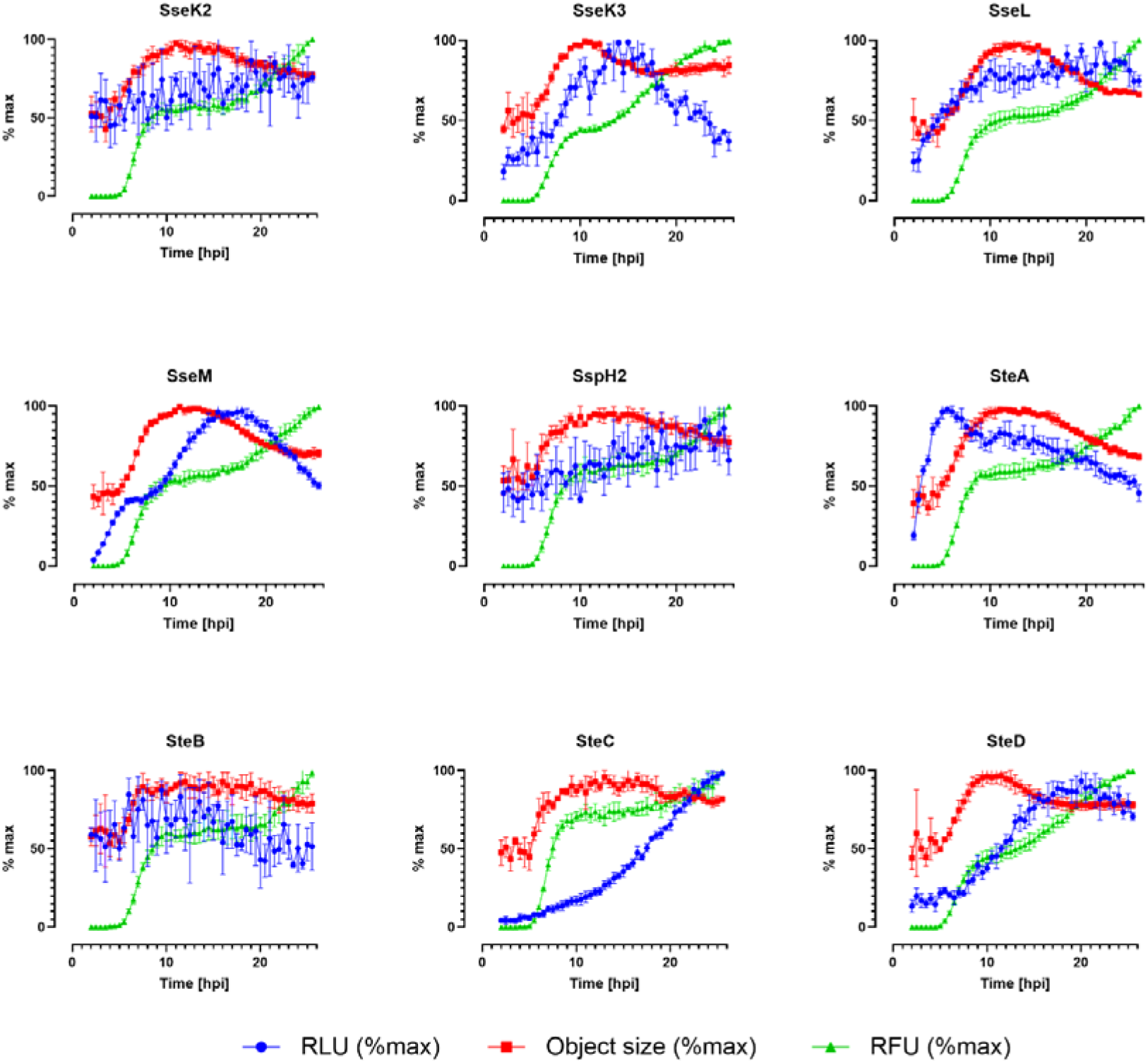
Real-time profiling of intracellular replication dynamics and NanoLuc complementation–based effector translocation of HiBiT-tagged *Salmonella* SL1344 strains in HeLa cells. HeLa cells stably expressing LgBiT were infected with a panel of 39 endogenously HiBiT-tagged Salmonella enterica serovar Typhimurium SL1344 strains, all generated in a constitutively eGFP-expressing background (*hisG*46 P_tet_::*gfp*; PVD0002). Infections were performed at a multiplicity of infection (MOI) of 50, and real-time monitoring was conducted for 24 hours (from 2 to 26 hpi) using automated multimodal imaging in a plate reader under environmental control. Total green fluorescence (RFU, green) and average green object size (µm^2^, red) serve as proxies for intracellular bacterial load and bacterial distribution within infected cells, respectively. Luminescence signal (RLU, blue) reflects NanoLuc complementation following translocation of HiBiT-tagged effectors into the host cell cytosol. Fluorescence and luminescence parameters were normalized to their respective maxima (% max) to facilitate comparison across strains. Data points represent mean ± SD of technical quadruplicates. Individual strain identifiers and effector annotations are listed in **Supplementary Table S2**.

**Supplementary Figure S4.**
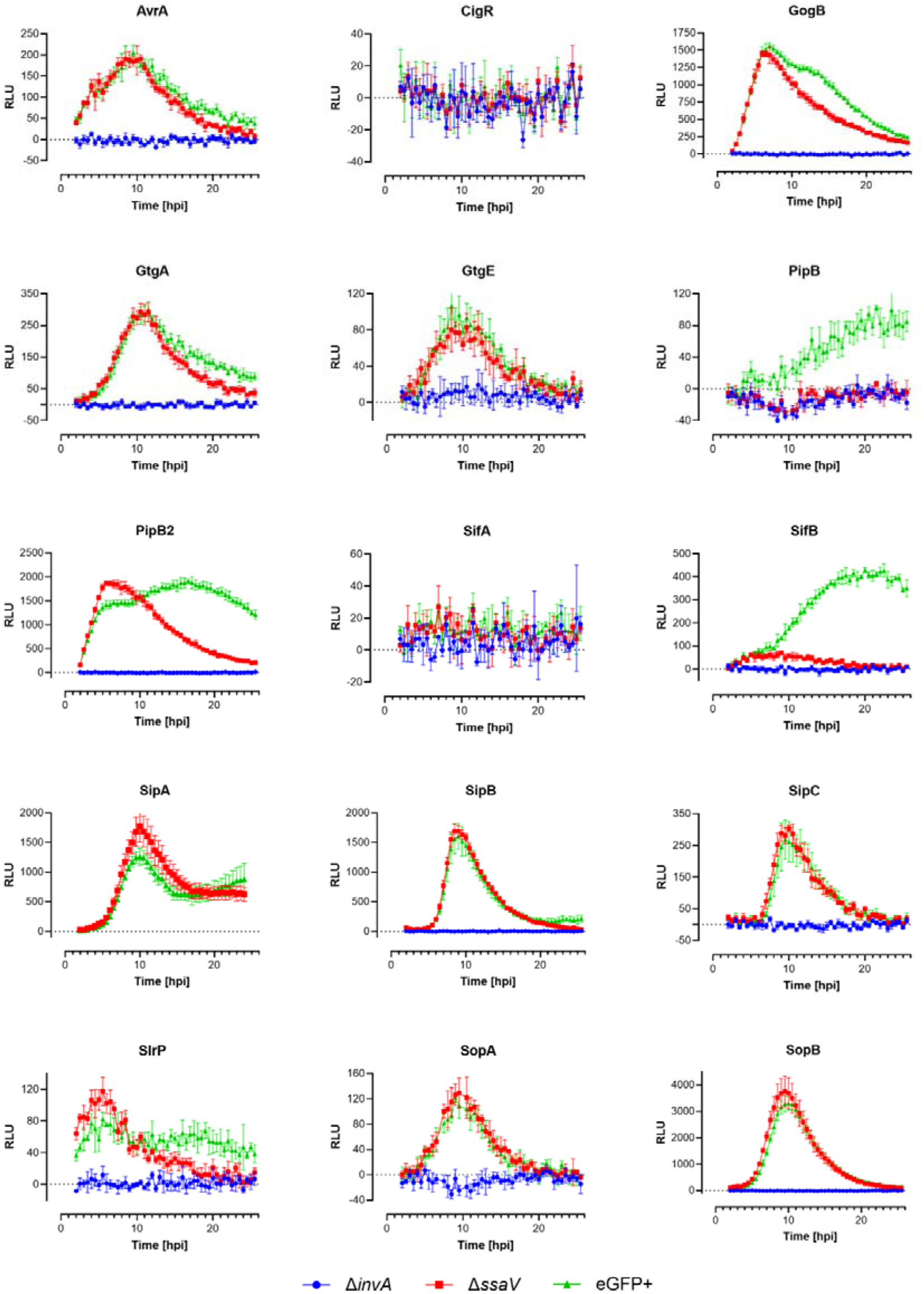
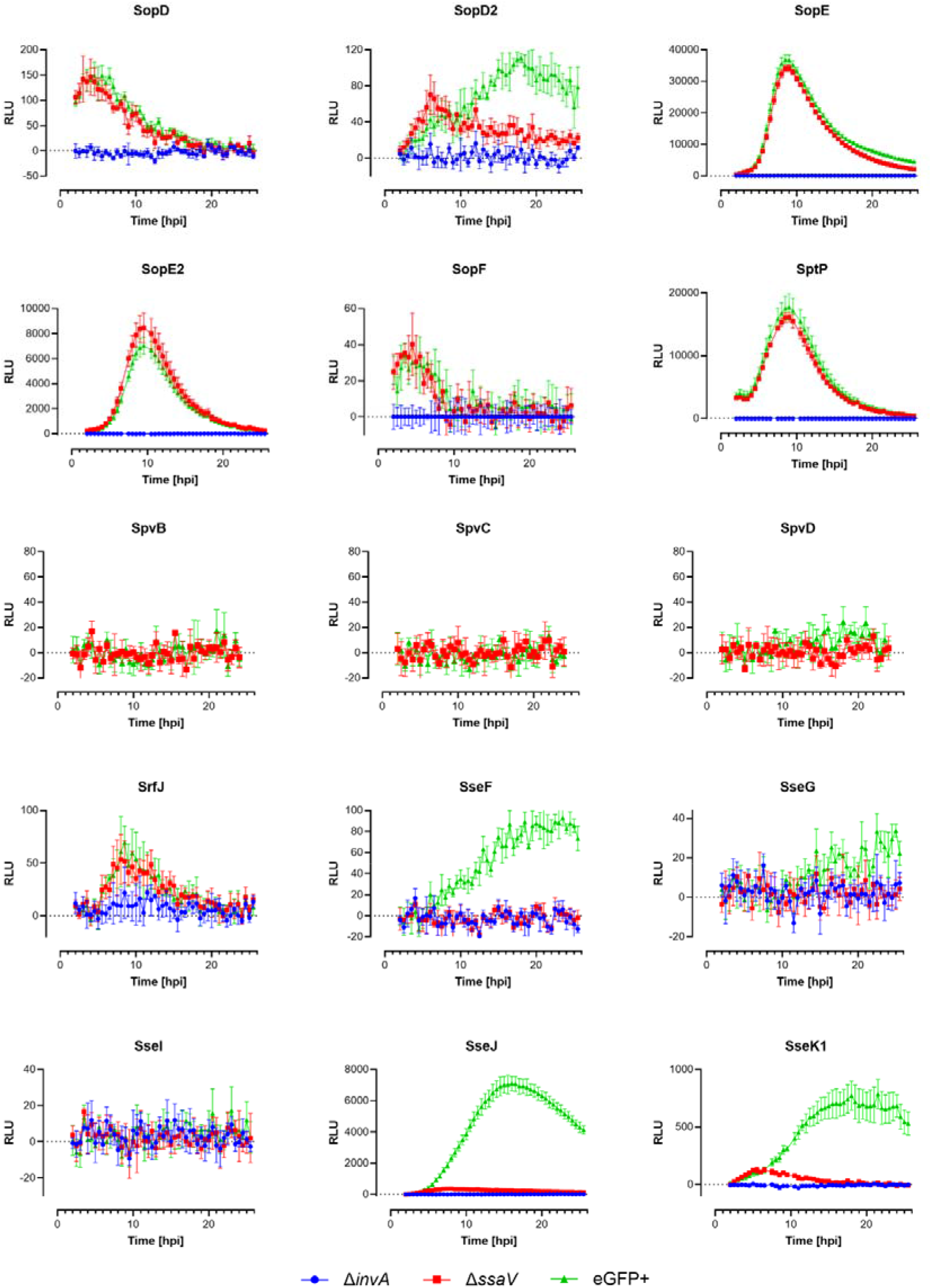
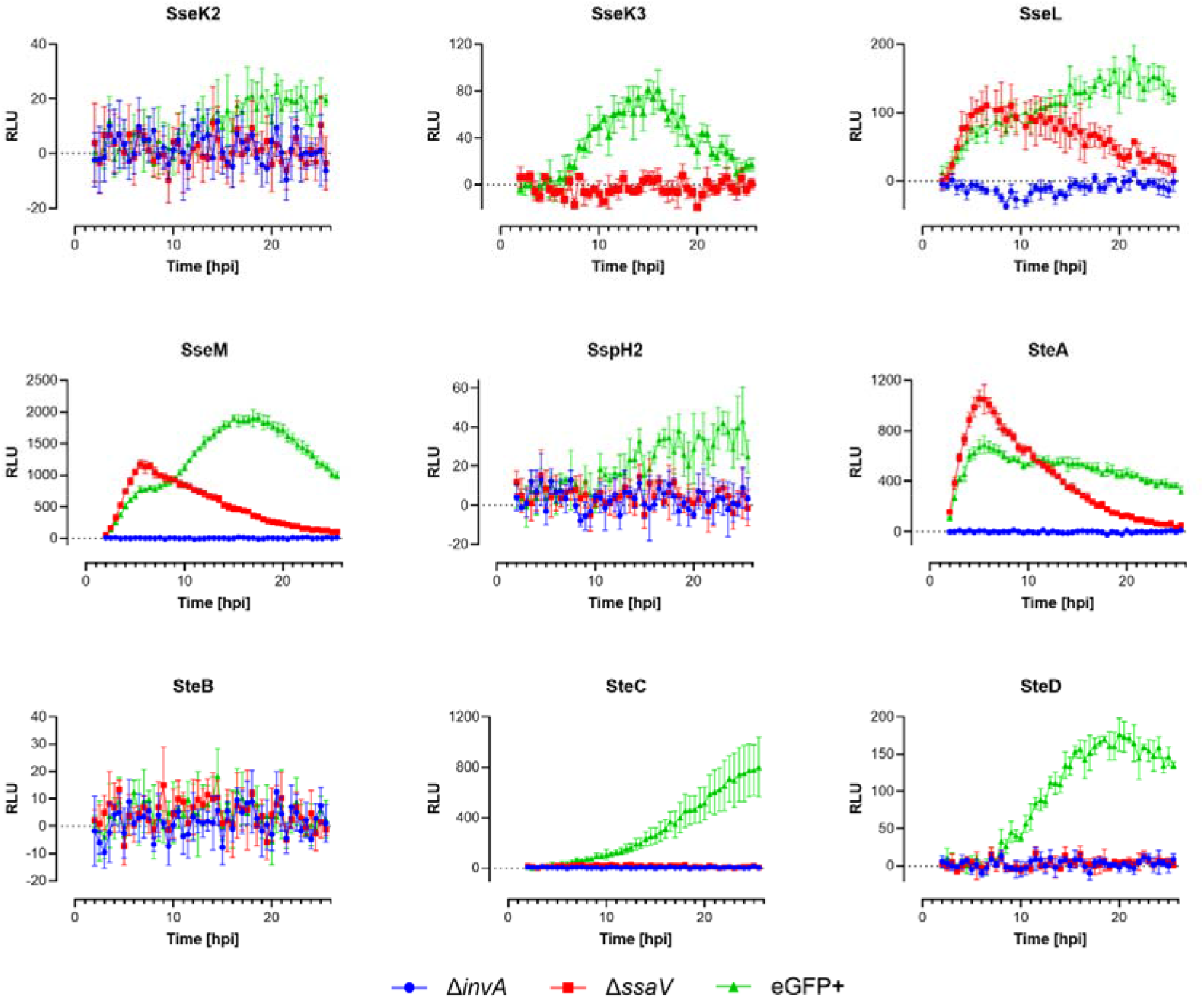
Real-time NanoLuc-based profiling of effector translocation dynamics across the *Salmonella* type III effector repertoire in wild-type and secretion system-deficient backgrounds during epithelial infection. HeLa cells stably expressing LgBiT were infected with endogenously HiBiT-tagged Salmonella enterica serovar Typhimurium SL1344 effector strains in three genetic backgrounds: wild-type SL1344 carrying constitutive eGFP^+^ (*hisG*46 P_tet_ ::*gfp*; PVD0002; green), the T3SS-1–deficient *ΔinvA::cat* mutant (PVD0316; blue), and the T3SS-2–deficient *ΔssaV::cat* mutant (PVD0317; red). Luminescence profiles (RLU), reflecting NanoLuc complementation following effector translocation into the host cytosol, were recorded from 2 to 26 hours post-infection (hpi). Infections were performed at a multiplicity of infection (MOI) of 50, and real-time luminescence monitoring was conducted using a Spark Cyto 400 multimodal plate reader under environmental control in the presence of Nano-Glo Vivazine. Luminescence values were background-subtracted (non-infected HeLa). Effectors are displayed alphabetically and distributed across panels (A–C). Data points represent mean ± SD of technical quadruplicates. Individual strain identifiers and effector annotations are provided in **Supplementary Table S2**.

**Supplementary Figure S5.**
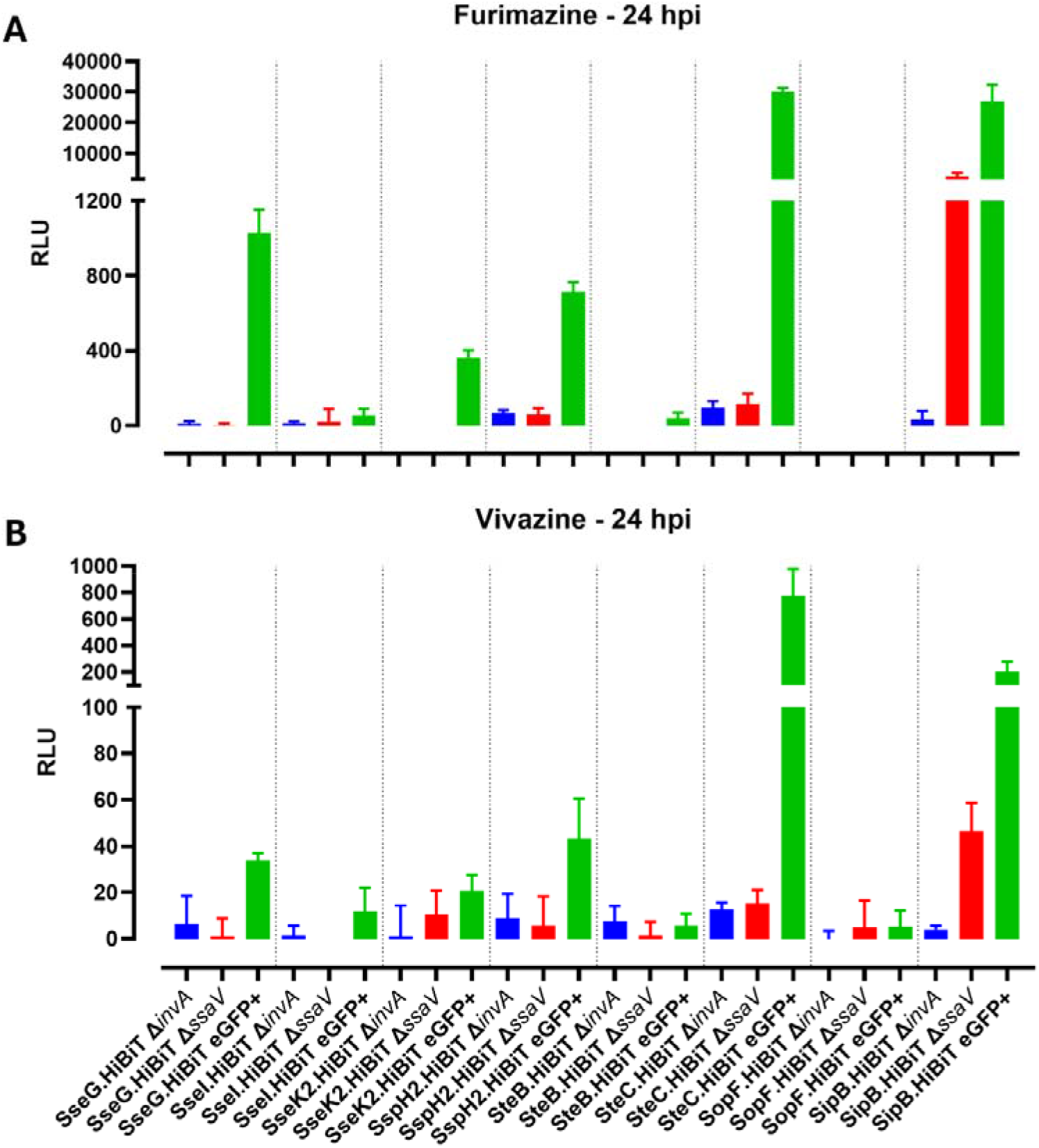
Comparative luminescence profiling of selected *Salmonella* type III effectors using vivazine- and furimazine-based NanoLuc complementation at 24 hours post-infection. HeLa cells stably expressing cytosolic LgBiT were infected with selected HiBiT-tagged S. Typhimurium SL1344 strains in a constitutively eGFP-expressing background (*his*G46 P_tet_::gfp; PVD0002), including T3SS-1–deficient (*ΔinvA;* blue) and T3SS-2–deficient (*ΔssaV;* red) derivatives. Luminescence was measured at 24 hours post-infection (hpi), corresponding to the indicated timepoint from continuous real-time acquisition using (A) furimazine, added shortly before readout, or (B) vivazine, present throughout infection. Bars represent raw luminescence values (RLU; mean ± SD of technical replicates). Strain identifiers and effector annotations are listed in **Supplementary Table S2**.

**Supplementary Figure S6.**
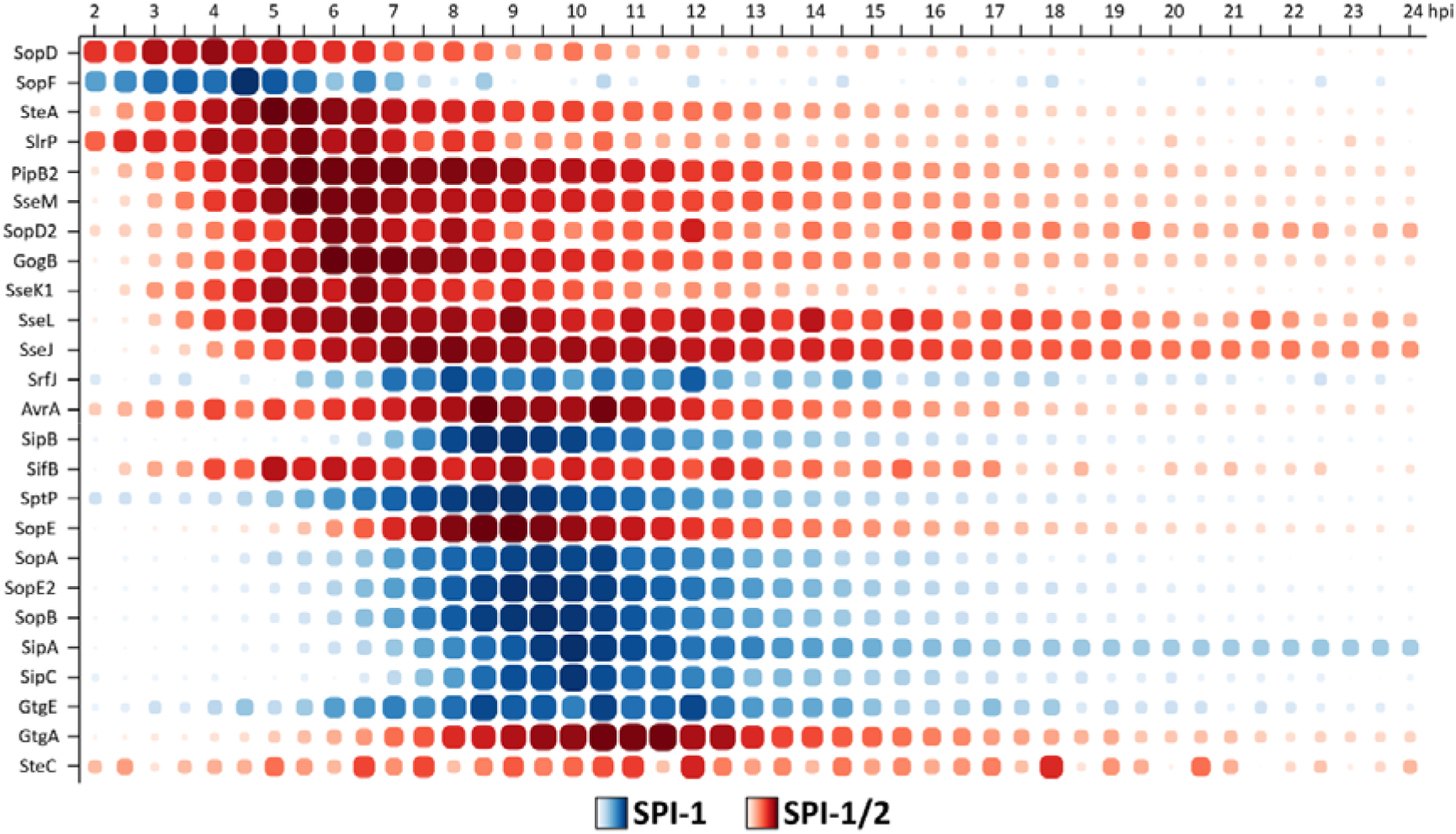
Hierarchical organization of effector translocation timing in the absence of T3SS-2. Matrix plots depict the temporal progression of effector translocation in the ΔssaV background, based on real-time HiBiT luminescence measurements normalized to the maximal signal observed for each effector (% max). Values represent means of technical quadruplicates. Effectors are ordered vertically according to ascending time of maximal translocation (T_max_), revealing the hierarchy of SPI-1–dependent effector delivery when T3SS-2 function is absent. Only effectors that met the criteria for detectable translocation in this background are shown (25 effectors). Each column represents a time point from 2 to 24 hours post-infection (hpi). Dot size and color intensity indicate the relative proportion of maximal translocation at each time point (% max). Effectors are color-coded by secretion system dependence as determined by phase-resolved AUC analysis: SPI-1–dependent (blue), and dual SPI-1/SPI-2–dependent (red). For dual-system substrates, the relative contribution of SPI-2 to total cumulative translocation (calculated from WT vs. ΔssaV AUC values) is indicated on the right-hand axis as percentage of SPI-2–dependent delivery. Across the ΔssaV dataset, SPI-1– dependent effectors exhibited a median T_max_ of 9.5 hpi, whereas the SPI-1 component of dual-system effectors peaked earlier, with a median T_max_ of 6.5 hpi, indicating preferential early deployment of dual substrates during intracellular SPI-1 activity in the absence of SPI-2–mediated secretion.

## SUPPLEMENTARY TABLES

**Supplementary Table S1. Annotated type III effector repertoire encoded by Salmonella enterica serovar Typhimurium SL1344.** Overview of the 39 currently identified type III effector (T3E) proteins encoded by *Salmonella enterica* serovar Typhimurium strain SL1344. For each effector, information is listed on gene identity with full and alternative names; SPI expression/secretion assignment (SPI-1, SPI-2, dual SPI-1/SPI-2, or N.D. when not determined or not conclusively resolved)^2, 20^; indication of SPI-1 translocon components, protein sequence length, and predicted molecular weight (with and without C-terminal HiBiT tag); genomic locus identifiers; UniProt and GenBank accession numbers; and reported or putative biochemical functions. Previously reported secretion and/or translocation outcomes obtained using C-terminally tagged effector variants (either plasmid-encoded or chromosomally integrated) are included where available, alongside tag identity, the experimental infection model used to evaluate translocation, and a corresponding publication reference (PMID). Where secretion or translocation status is supported only by indirect or in vitro evidence, this is marked by an asterisk (*). Abbreviations: SPI, *Salmonella* pathogenicity island; T3SS, type III secretion system; N.D., not determined/not conclusively assigned; MW, molecular weight; aa, amino acids; N/A, no published information available.

**Supplementary Table S2. Strain collection of endogenously HiBiT-tagged type III effectors in Salmonella enterica serovar Typhimurium SL1344**. For each of the 39 annotated type III effector proteins encoded by SL1344, the corresponding in-house strain identifiers (PVDxxxx) are listed for endogenously HiBiT-tagged derivatives constructed in four genetic backgrounds: wild-type (WT) SL1344 (*hisG*46; PVD0001), GFP-expressing SL1344 (*hisG*46 P_tet_::*gfp*; PVD0002), SL1344 Δ*invA*::*cat* (PVD0316), and SL1344 Δ*ssaV*::*cat* (PVD0317). All strains were generated by λ Red–mediated recombineering and encode C-terminal HiBiT fusions at the native genomic locus of the respective effector gene. The helper plasmid used for recombineering is indicated for each effector. Effectors are listed in alphabetical order. Background strains used for strain construction are described in Supplementary Table S3. Abbreviations: *HiBiT*, 33 bp tag-encoding sequence coding for an 11-amino-acid luminescent peptide tag; *aphT*, kanamycin resistance gene; *cat*, chloramphenicol resistance gene; FRT, flippase (FLP) recombinase recognition target site.

**Supplementary Table S3. Background Salmonella enterica serovar Typhimurium SL1344 strains used for HiBiT-tagged type III effector strain construction.** Overview of the SL1344 background strains used for the generation of endogenously HiBiT-tagged effector derivatives. In-house strain identifiers (PVDxxxx) are listed together with the corresponding genotypes, including the genomic loci targeted by insertions or deletions, and references to the original strain sources where applicable. Deletion background strains were generated by λ Red–mediated recombineering using pKD46 as the helper plasmid. Abbreviations: *aphT*, kanamycin resistance gene; *cat*, chloramphenicol resistance gene; FRT, flippase (FLP) recombinase recognition target site; tko, translational knockout.

**Supplementary Table S4. Oligonucleotide primers used for λ Red recombineering and sequence verification in *Salmonella enterica* serovar Typhimurium SL1344**. Overview of oligonucleotide primers used in this study for endogenous C-terminal HiBiT-tagging of effector genes, targeted gene deletions and mutations, allelic replacements, and sequence verification of modified genomic loci. For each primer, the target locus, primer orientation (forward or reverse), and the intended genomic modification are indicated. Primers used for λ Red recombineering were designed to generate PCR-derived linear editing substrates carrying 50 bp homology arms flanking the targeted genomic region (capital letters). For HiBiT-tagging, forward primers encode the 33-bp *HiBiT* sequence (underlined sequences) immediately upstream of the native stop codon of the effector gene. For gene deletions, primers amplify antibiotic resistance cassettes flanked by homology regions corresponding to either the full coding sequence or defined internal regions in the case of non-polar deletions. Resistance cassettes were derived from pKD3 (*cat*) or pKD4 (*aphT*), in some cases flanked by FRT sites. Single-stranded DNA oligonucleotides for allelic replacement (i.e., translational knockout) were designed with the Mage Oligo Design Tool (MODEST)^67^. Lowercase letters indicate nucleotides targeted for mutation. Sequencing primers used to verify correct genomic modification are included. Targeted gene loci are listed alphabetically. Abbreviations: *HiBiT*, 33 bp tag-encoding sequence coding for an 11-amino-acid luminescent peptide tag; *aphT*, kanamycin resistance gene; *cat*, chloramphenicol resistance gene; FRT, flippase (FLP) recombinase recognition target site; FW, forward primer; REV, reverse primer; nt, nucleotides; OMAR, oligo-mediated allelic replacement.

**Supplementary Table S5. Quantitative summary of effector translocation kinetics and secretion system dependence.** For each HiBiT-tagged effector, luminescence time-course data were quantified in wild-type (WT), Δ*ssaV* (T3SS-2–deficient), and Δ*invA* (T3SS-1–deficient) backgrounds using real-time NanoLuc complementation assays. Area-under-the-curve (AUC; expressed as relative light units × hours, RLU·h) values were calculated by trapezoidal numerical integration after background subtraction over defined infection intervals. SPI dependence (this study) indicates classification as SPI-1–dependent, SPI-2–dependent, or dual SPI-1/SPI-2–dependent based on phase-specific statistical comparisons of WT, Δ*invA*, and Δ*ssaV* strains (see Methods). SPI dependence (literature) summarizes previously reported secretion system assignments (see also **Supplementary Table S1**). SPI-1 detectable and SPI-2 detectable indicate whether statistically significant translocation was detected. SPI-1 and SPI-2 contributions (expressed as percentages of the WT AUC) were calculated for effectors showing detectable WT translocation as follows: SPI-1 contribution (%) = (Δ*ssaV* AUC / WT AUC) x 100); SPI-2 contribution (%) = 100 - SPI-1 contribution (%). WT mean AUC (2–24 hpi), Δ*ssaV* mean AUC (2–24 hpi), and Δ*invA* mean AUC (2–24 hpi) report cumulative translocation (if any) over the full infection course. Max mean signal indicates the highest mean luminescence value observed during the time course. T_max_, T_50_, and T_25_ represent the first time point at which the mean signal reached the maximum, ≥50% of maximum, or ≥25% of maximum, respectively, providing measures of delivery onset and peak timing. WT mean AUC (2–10 hpi) and WT mean AUC (10–24 hpi) quantify mid-stage and late-stage translocation contributions. Percentage columns report the fraction of total effector signal contributed by each effector within the indicated phase or over the full infection period. Peak window start/end (≥90% max) define the continuous time interval during which translocation remained at or above 90% of maximal signal, providing an estimate of sustained peak delivery duration. Values represent means of technical quadruplicates.

